# *Trans*-eQTL mapping in gene sets identifies network effects of genetic variants

**DOI:** 10.1101/2022.11.11.516189

**Authors:** Lili Wang, Nikita Babushkin, Zhonghua Liu, Xuanyao Liu

## Abstract

Nearly all trait-associated variants identified in GWAS are non-coding. The *cis* regulatory effects of these variants have been extensively characterized, but how they impact gene regulation in *trans* has been the subject of much fewer studies. Mapping *trans* genetic effects is very challenging because their effect sizes tend to be small and a large multiple testing burden reduces the power to detect them. In addition, read mapping biases can lead to many false positives. To reduce mapping biases and substantially improve power to map *trans*-eQTLs, we developed a pipeline called trans-PCO, which combines careful read and gene filters with a principal component (PC)-based multivariate association test. Our simulations demonstrate that trans-PCO substantially outperforms existing *trans*-eQTL mapping methods, including univariate and primary PC-based methods. We applied trans-PCO to two gene expression datasets from whole blood, DGN (*N* = 913) and eQTLGen (*N* = 31,684), to identify *trans*-eQTLs associated with gene co-expression networks and hallmark gene sets representing well-defined biological processes. In total, we identified 14,985 high-quality *trans*-eSNPs–module pairs associated with 197 co-expression gene modules and biological processes. To better understand the effects of trait-associated variants on gene regulatory networks, we performed colocalization analyses between GWAS loci of 46 complex traits and *trans*-eQTLs identified in DGN. We highlight several examples where our map of *trans* effects helps us understand how trait-associated variants impact gene regulatory networks and biological pathways. For example, we found that a locus associated with platelet traits near *ARHGEF3 trans-*regulates a set of co-expressed genes significantly enriched in the platelet activation pathway. Additionally, six red blood cell trait-associated loci *trans*-regulate a gene set representing heme metabolism, a crucial process in erythropoiesis. In conclusion, trans-PCO is a powerful and reliable tool that detects *trans* regulators of cellular pathways and networks, which opens up new opportunities to learn the impact of trait-associated loci on gene regulatory networks.

## Main

Genome-wide association studies (GWAS) have identified tens of thousands of genetic loci associated with a large number of complex traits and diseases. More than 90% of GWAS loci are located in non-coding regions of the genome and are thought to affect human traits by regulating gene expression^1–5^. Nearly all studies to date have focused on understanding the effects of trait-associated variants on gene expression in *cis*, which only include effects on genes that are near the associated loci. However, multiple lines of evidence suggest *cis-* regulatory effects only capture a small proportion of the heritability of complex traits and diseases. For example, Yao et al.^6^ estimated that only an average of 11% of trait heritability is explained by *cis*-genetic effects on gene expression levels. We previously hypothesized that *trans*-eQTLs, despite having very small effects on each individual gene, may cumulatively account for a large proportion of trait variance^7^. Indeed, our modeling indicates that *trans*-eQTL effects account for twice as much genetic variance in complex traits as *cis*-eQTL effects^7^. Thus, establishing a representative map of genetic variants and their *trans* effects is a critical step toward understanding complex trait and disease genetics.

Two major challenges have precluded *trans*-eQTL discovery to date. First, *trans*-eQTL mapping is extremely prone to false positives due to mapping errors that cause short sequences to map to homologous regions of the genome^8^. This causes spurious associations between the mapping coverages at multiple homologous regions, which result in strong but artificial *trans*-eQTL signals if unaccounted for. The second challenge is by far more difficult to overcome: *trans*-eQTLs are challenging to detect compared to *cis*-eQTLs because (i) they have much smaller effect sizes than *cis*-eQTLs^7^, and (ii) a genome-wide search of *trans*-eQTLs involves a huge number of statistical tests (∼10^6^ SNPsx ∼20k genes) resulting in a heavy burden of multiple testing corrections.

Previous work in yeast and human cells suggests that *trans*-eQTLs generally affect the expression levels of multiple genes. In particular, Albert et al.^9^ found that the 90% of *trans*-eQTLs in their yeast segregant system could be mapped to just 102 hotspot loci that regulate a median of 425 genes each. In fact, three hotspots were found to affect over half of the 5600 expressed *trans*-regulated genes, indicating that some *trans*-eQTLs may have significant genome-wide effects. In humans, the largest *trans*-eQTL study to date from eQTLGen^10^ (n = 31,684) identified 59,786 *trans*-eQTL signals for 3,853 SNPs, indicating that each locus affects at least 15 genes on average. Thus, compared to the traditional approach of testing the association between genetic variants and the expression level of a single gene^11,12^, testing *trans* associations between genetic variants and the expression levels of a group of genes can greatly improve the power to identify *trans*-eQTLs.

Indeed, many disease-associated loci are “peripheral master regulators”, which regulate multiple genes in the core disease-relevant pathways^7,13^. For example, *KLF14*, which is significantly associated with type 2 diabetes, is a peripheral master regulator that modulates the expression of 385 genes involved in lipid metabolism^14^. Other examples include the *FTO/IRX3/IRX5* locus in obesity^15^ or the p53 tumor-suppressor gene in cancers^16^. A method that can detect a large number of *trans*-eQTLs associated with multiple genes in gene networks would allow functional interpretation of more disease associated loci and shed light on the underlying mechanisms.

The co-regulation and co-expression patterns of genes driven by *trans*-eQTL have long been recognized. Yet, most methods do not directly map *trans*-eQTLs of co-expressed gene sets, but rather use the coexpression patterns to improve *cis*-eQTL or *trans*-eQTL mapping of a single gene. Some utilized the co-expression patterns of genes to account for hidden variation in *trans*-eQTL analysis, and thus improves power while reducing false *trans*-eQTL discovery. For example, Joo et al.^17^ used the global correlation structure of genes to capture and remove confounding effects from *trans* associations. Similarly, Rakitsch and Stegle^18^ utilized the local co-expression patterns of *trans*-target genes to infer the appropriate covariates to be included in *trans*-eQTL association testing. Zhou and Cai^19^ jointly modeled the effects of *cis* regulatory variants and gene regulatory networks on expression levels of a target gene, therefore allowing the simultaneous identification of *cis*-eQTL and regulatory networks; the model then identifies individual *trans*-eGenes, which are mediated by the *cis* regulatory effects. Several studies aimed to identify trans-eQTLs of co-expressed genes. For example, some studies^20,21^ used tensor decomposition to decompose gene expressions of multiple tissues into a few latent components, which might capture gene co-expression, to identify *trans*-eQTLs of the components. However, the method requires gene expression data of multiple tissues, which is not readily available for many gene expression studies. Rotival et al.^22^ used independent component analyses to identify co-expression gene sets, and subsequently tested for the enrichment of *trans* signals in the gene sets by hypergeometric tests. More recently, Kolberg et al.^23^ identified 38 *trans*-eQTLs (10% FDR) associated co-expressed gene modules, by testing the association between SNPs and a single ‘eigengene’ (essentially the primary principal component, PC1) of gene modules that captures the co-expression pattern. Nonetheless, PC1 has very limited power at identifying genetic effects (see below and Figure 2). Dutta et al.^24^ leveraged canonical correlation analysis to identify *trans* associations between multiple disease–associated SNPs and multiple genes, by integrating with GWAS signals. However, the method has different goals than identifying *trans*-eQTLs of multiple genes in specific tissues (for example, it is useful for identifying “core” like disease genes and processes for a specific disease; see Discussion and Supplementary Note).

Our main goal is to develop a method for detecting *trans*-eQTLs associated with multiple genes in a gene module by using multivariate association. Multivariate association methods tend to be more powerful than univariate association methods. Detecting *trans*-eQTLs of gene modules containing multiple co-regulated genes can also potentially improve power by reducing multiple testing burdens, because the number of tested gene modules is much less than the number of genes. However, there are caveats. First, sequence similarity among distinct genomic regions can lead to severe false positive discovery issues in trans-eQTL mapping^8^. This is especially problematic in mapping trans-eQTLs of co-expression gene modules because genes can be falsed clustered due to sequence similarities^8^. For example, the top latent components in study^20^ mostly represent genes sharing homologous sequences. Therefore, we need to be extra diligent about multi-mapping of sequencing reads when mapping *trans*-eQTLs of co-expression gene modules. Second, the naïve way of using a single component, such as the first gene expression PC, to represent the gene modules and use it as the response variable in association tests, can significantly reduce power. While the primary PC captures the largest amount of total variance in gene expression levels, it can be less powerful or even powerless in detecting significant associations than higher-order PCs, because the direction of the genetic effects on the genes may not align with the primary PC^25,26^. It is also difficult to predict which higher-order PC has the highest power^26^.

To combat this, we propose trans-PCO, a flexible approach that uses the PCA-based omnibus test^26^ (PCO) to combine multiple PCs and improve power to detect *trans*-eQTLs. Trans-PCO tackles both major challenges in *trans*-eQTL mapping. First, trans-PCO carefully filters sequencing reads and genes based on mappability across different regions of the genome to avoid false positives due to multi-mapping^8,27,28^. Second, trans-PCO uses a novel multivariate association test^26^ to detect genetic variants with effects on multiple genes in predefined sets and captures genetic effects on multiple PCs. By default, trans-PCO defines sets of genes based on co-expression gene modules as identified by WGCNA^29^. It also accepts user-defined sets; for example, genes that belong to the same gene ontology^30^, KEGG pathway^31^, or protein complex^32^.

We applied trans-PCO to RNA-sequencing data from the Depression Genes and Networks study^27^ (DGN, sample size *N* = 913) and the summary-level statistics from the eQTLGen study^10^ (sample size *N* = 31,684) to identify *trans*-eQTLs associated with co-expression gene modules and well-defined biological processes in whole blood. In total, trans-PCO identified 14,985 high-quality *trans*-eSNPs–module pairs associated with 197 co-expression gene modules and biological processes. We also performed colocalization analysis^33^ of GWAS loci of 46 complex traits and *trans*-eQTLs, in order to explore how trait-associated variants impact gene regulatory networks and pathways in *trans*. All *trans*-eQTLs that are associated with gene co-expression networks and biological pathways can be found in www.networks-liulab.org/transPCO.

## Results

### Overview of the method

We developed the trans-PCO pipeline to detect *trans*-eQTLs that are associated with the expression levels of a group of genes (gene module) by using a PC-based multivariate association test^26^ that combines multiple gene expression PCs. The trans*-*PCO method consists of three main steps (Figure 1).

First, trans-PCO pre-processes RNA-seq data to reduce false positive *trans*-eQTL associations due to read multimapping errors. Specifically, trans-PCO removes all sequencing reads mapped to low mappability regions of the genome (mappability score <1, Methods) before profiling gene expression levels. These procedures substantially reduce the occurrence of false positive *trans*-eQTLs due to sequencing alignment errors^8,28^. When only summary-level data are available (e.g., eQTLGen dataset^10^), trans-PCO dynamically excludes from the module any genes that are cross-mappable to genes within 100 kb of the tested SNP.

Second, trans-PCO groups genes into clusters, which alleviates the burden of multiple testing by reducing the number of statistical tests and thus increases the statistical power. By default, trans-PCO determines the gene groupings by using WGCNA^29^ to identify co-expression modules from gene expression levels (see Methods). We remove covariates and confounders, such as batch effects, gene expression PCs, and cell type proportions (e.g., DGN dataset^27^), from gene expression levels before grouping gene modules. This step is necessary to ensure that the gene modules are not primarily driven by confounding factors. Trans-PCO also allows customization of the gene groups or sets, i.e., genes in the same pathway or protein-protein interaction network^30–32^ can be grouped into user-defined gene modules.

Lastly, trans-PCO tests for association between each SNP and the expression levels of the genes in each gene module by adapting the PCO method, which combines multiple gene expression PCs by using six PC-based statistical tests: PCMinP, PCFisher, PCLC, WI, Wald and VC (see Methods for details). Each PC-based test combines multiple PCs uniquely, which allow signals under various genetic architectures to be captured. PCO evaluates the six PC-based tests and takes the minimum p-value as the final test statistic. The final p-values are computed according to Liu et al.^26^ (also see Methods). Only PCs with eigenvalues λ_k_> 0.1 are used in trans-PCO (See Supplementary Note). To avoid identifying associations driven by *cis*-effects, we excluded from the module all genes on the same chromosome as the test SNP. To correct for multiple testing, we performed 10 permutations to establish an empirical null distribution of p-values (See Supplementary Note).

**Figure 1.**
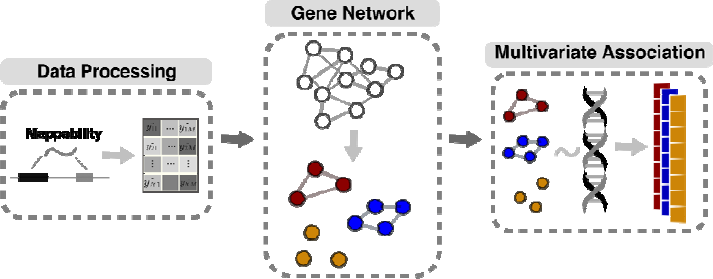
Three main steps in trans-PCO pipeline. The first step of trans-PCO pre-processes RNA-seq data to reduce false positive *trans*-eQTL associations due to read alignment errors. The second step involves grouping genes into gene sets, such as co-expression modules or biological pathways. The last step tests for *trans*-eQTLs of each gene set by a PC-based multivariate association test^17^.

### Trans-PCO outperforms existing methods in simulations

We performed simulations to evaluate the power of trans-PCO in detecting *trans*-eQTLs associated with multiple genes. We primarily compared the power to (i) the standard univariate test (“MinP”), and (ii) the primary PC-based test (“PC1”, Kolberg et al.^23^; see Methods). We used a co-expression gene module consisting of 101 genes from the DGN dataset (module 29). In null simulations, we simulated the z-scores between a SNP and *K* = 101 genes in a gene module following the null distribution, *N_K_* (0, *Σ_K×K_*), where *Σ_K×K_* is the residualized expression correlation matrix. In power simulations, we generated z-scores from the distribution 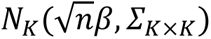 , where *n* is sample size (*N* = 200,400,600, and 800) and *β* is a vector representing the true effect sizes of the SNP on *K* genes. Among *K* genes, a proportion *γ* of them are causal with non-zero effects. Therefore, we generated *β_k_* from a point normal distribution, where *β_k_* ~ *N* (0, *σ*^2^*b*) for proportion *γ* (*γ* = 1%, 5%, 10%, 30% *and* 50%), and *β_k_* = 0, otherwise. The *trans-*genetic variance is *σ*^2^*b* (*σ*^2^*b* = 0.001), which is a low and realistic per SNP heritability for *trans* effects. We simulated 10,000 SNPs and performed 1000 simulations.

To compare the power of different approaches to identify *trans*-eQTLs, we defined significant univariate tests for a gene module based on the minimum p-values across *K* genes. For the primary PC-based method, we used PC1 of the gene module co-expression matrix to represent the module and tested it for associations. In power simulations, we set significance levels at 10% FDR to be consistent with real data analysis. We computed power as the average proportion of significant tests out of 10,000 simulated SNPs across 1000 simulations.

Trans-PCO significantly outperformed the univariate test and the primary PC method across different sample sizes and proportions of causal genes **(Figure 2)**. Specifically, the power of trans-PCO increases rapidly with increasing sample sizes. At the sample size of 800, assuming 30% of genes have causal effects in the gene module, the power of trans-PCO is 74%, compared to 15% for the univariate test and 0.0018% for the primary PC method **(Figure 2A)**.

We also compared the power of each method across various causal gene proportions using a fixed sample size (n=500). All three methods have little power in detecting *trans*-eQTLs when the proportion of causal genes is below 10%. However, above this threshold, the power of trans*-* PCO increases dramatically: 36% at 30% causal genes and 86% at 50% causal genes. In contrast, the univariate and the primary PC methods remain almost powerless for nearly all simulated scenarios (**Figure 2B**). We note that the primary PC method appears to be almost powerless across the scenarios, which agrees with the previous observation that the primary PC can be less powerful than higher-order PCs in GWAS^11^. Simulation results at various genetic variances can be found in Supplementary Materials, including at extremely low proportions of causal genes and high *trans* effects (Figure S1). We found that the univariate method only outperforms trans-PCO when the proportion of causal genes is extremely low, such as only one causal gene in the entire gene set, and the *trans* effects are large. Trans-PCO gains more power when there are more than 1 causal gene, as it aggregates multiple weak effects to improve power. Null simulations demonstrated all three methods are well-controlled for false-positive inflations (Figure S2).

**Figure 2.**
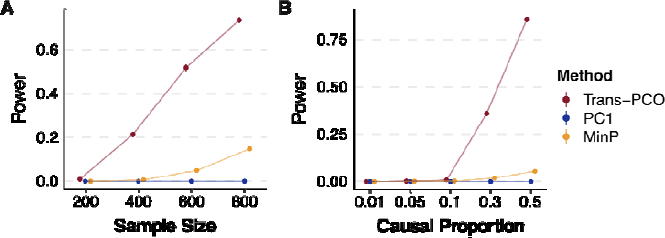
Power of trans-PCO across different sample sizes and causal gene proportions, in comparison to PC1 (Kolberg et al.^23^) and univariate (MinP) methods. **(A) Power comparison across various sample sizes (*N* = 200, 400, 600, and 800)**. *Trans-*genetic variance was simulated to be 0.001 and the proportion of causal genes in the gene module was 30%. Error bars representing 95% confidence intervals are plotted, but many are too small to be visible. See numerical results in Table S2. **(B) Power comparison across different proportions of causal genes in the gene module.** The simulated sample size was 500. Points show average power across 1000 simulations. Error bars representing 95% confidence intervals are plotted, but many are too small to be visible. See numerical results in Table S2.

We also included comparisons to two additional methods: ARCHIE proposed by Dutta et al.^24^ and a method by Rovital et al.^22^ (see Supplementary Note, Figure S19 and Figure S20). We showed that ARCHIE is not powerful at detecting *trans*-eQTL effects from a SNP to multiple genes, which are the effects trans-PCO was designed for (Figure S19D). We note that the main goal of ARCHIE is to identify trait-specific gene sets associated with GWAS loci, whereas trans-PCO is designed to map *trans*-eQTLs for any user-specified gene sets in specific tissues or cell types (see Supplementary Note, Figure S19 and Discussion). Rovital et al.^22^ is based on the primary PC-based approach and we showed that the method also has limited power at identifying weak *trans*-eQTL effects (Supplementary Note and Figure S20).

### Trans-PCO identifies 3899 *trans*-eSNP–module pairs associated with co-expression gene modules in the DGN dataset

We used trans-PCO to identify *trans*-eQTLs associated with co-expression gene modules in RNA-seq data from whole blood samples of the DGN cohort (*N* = 913)^27^. WGCNA^29^ identified 166 co-expression gene modules, with the number of genes in each module ranging between 625 (module 1, M1) and 10 (module 166, M166) (Supplementary Table S1). We then performed genome-wide scans of *trans*-eQTLs for each gene module. At 10% FDR, trans-PCO identified significant *trans*-eQTLs for 102 out of 166 gene modules, corresponding to 3899 significant *trans*-eSNP–module pairs (Supplementary Table S3). Many trans-eSNPs are in linkage-disequilibrium (LD). Using LD clumping to group *trans*-eSNPs into LD-independent loci (R2<0.2), we found 202 *trans*-loci–module pairs (**Figure 3A**, Table S3, Table S4).

We compared *trans*-eQTL signals detected in DGN by trans-PCO to signals identified by the univariate method in Battle et al.^27^. Out of 12,132 genes analyzed by trans-PCO, the univariate method detected 326 significant *trans*-eSNP–gene pairs for 128 genes at 5% FDR^27^. At the same FDR level, trans-PCO identified 3031 significant *trans*-eSNP–gene module pairs for 75 gene modules. We compared the magnitude of the significant *trans*-eQTL effects detected by trans-PCO and the univariate method. More specifically, we compared the maximum univariate z-scores of SNPs and each gene in significant *trans*-eSNP–module pairs identified by trans-PCO to the z-scores of significant *trans*-eSNP–gene pairs by the univariate method. We found that the maximum z-scores of trans-PCO signals are much smaller than z-scores of the univariate method signals (**Figure 3B**), indicating that our multivariate approach can detect much smaller *trans* effects than univariate methods.

We also applied the primary PC method (Kolberg et al.^23^) to DGN, and identified 1483 significant *trans*-eSNP–module pairs (55 *trans*-loci–module pairs) at 10% FDR, and 1464 pairs (99%) were detected by trans-PCO (Figure S13A). Notably, in total, trans-PCO identified more than twice the signals than the primary PC method. However, the primary PC method identified more signals than expected, as it was previously shown to be powerless in the simulations. We note that we simulated weak effects and sparse causal proportions to better reflect common and realistic *trans* effects, and the primary PC method is powerless in these settings. We performed additional simulations with large effects and high causal proportions, and the primary PC method achieved 50% power as trans-PCO (see Supplementary Note, Figure S18). Additionally, we found in the DGN dataset that the univariate z-scores of *trans* signals detected by the primary PC method are larger than those of trans-PCO signals (Figure S13B-D). Therefore, the *trans* signals detected by the primary PC method are likely of strong *trans* effects, and trans-PCO is able to detect additional weak *trans* effects. Statistically, PC1 can also have good statistical power when the *trans* effects align with the direction of PC1^26^. To demonstrate this, we performed simulations under the assumption that the genetic effect vector perfectly aligns with the primary PC direction. We found that the primary PC method has higher power than trans-PCO and MinP methods under this special circumstance (Figure S14). However, it is impossible to predict when the genetic effects align with gene expression PCs.

### *Trans*-eQTLs are enriched in variants with *cis*-regulatory effects on transcription factors

We found that only 31 *trans*-eSNPs (1%) are in coding regions, suggesting that a very small proportion of *trans*-eQTLs impact gene expression levels in *trans* by altering protein coding sequences. Several studies have shown that *trans*-eQTLs have *cis*-regulatory effects, impacting the expression levels or splicing of nearby genes^10,27^; thus, we evaluated our identified *trans*-eQTLs for concomitant *cis*-regulatory activity. We first overlapped *trans*-eSNPs with *cis*-eQTLs and *cis*-splicing QTLs (*cis*-sQTLs) in DGN^34^. Of the 2955 *trans*-eSNPs (Table S3), we found that 71% are significant *cis*-eSNPs in DGN, and 46% are significant *cis*-sSNPs, together accounting for 73% of all *trans*-eSNPs. To further examine whether the *cis* and *trans* effects are driven by the same variant, we performed colocalization analysis of *trans*-eQTLs with *cis*-eQTLs and *cis*-sQTLs using coloc^33^ (see Methods). Specifically, we first grouped *trans*-eSNP–gene module pairs into 179 *trans*-region–gene module pairs, based on 200kb fixed-width regions (see Methods). We then performed colocalization analyses between the *trans*-eQTLs and *cis*-eQTLs/*cis*-sQTLs. We found that 51 out of 179 *trans* regions colocalized with a *cis*-eQTL (PP4>0.75, **Figure 3C** and Figure S3). 41 *trans*-loci colocalized with a *cis*-sQTL. Overall, 60 *trans*-loci share causal variants with at least one *cis*-eQTL or *cis*-sQTL (**Figure 3C**, Table S5), confirming that *trans*-eQTL effects are generally mediated through *cis*-gene regulation. Additionally, a large fraction of *trans*-loci (66%) do not colocalize with *cis*-eQTLs or *cis*-sQTLs. While power may have limited our ability to detect colocalization of some *trans*-eQTLs and *cis*-eQTLs, there might also exist unknown *trans*-regulatory mechanisms, independent of *cis* gene expression or splicing, which is subject to future studies.

We also investigated the types and functions of genes that are likely to mediate *trans*-eQTL effects. We found that the genes nearest *trans*-eQTLs are highly enriched in “RNA polymerase II transcription regulatory region sequence-specific DNA binding” (adjusted *P* = 1.26 x 10^-3^) and “DNA-binding transcription factor activity” (adjusted *P* = 1.39 x 10^-3^, Table S6), suggesting that transcription factors are important mediators of *trans*-eQTL effects. Indeed, trans-PCO identified and replicated several well-known master *trans* regulators in blood, such as *IKZF1*^28,35,36^, *NFKBIA*^28^*, NFE2*^10,28,37^, and *PLAGL1*^28,36^ (**Figure 3A**). We also found colocalization of these *trans*-eQTLs with *cis*-eQTLs at the *NFKBIA*, *NFE2* and *PLAGL1* loci (**Figure 3D**, Figure S3), supporting the conclusion that these genes are likely the *cis*-mediating genes.

### High quality map of *trans*-eSNP to gene module associations improves functional interpretation

Most of the gene modules used in trans-PCO have functional annotations, which allows us to interpret the functional roles of the *trans*-eQTLs identified by the method. We first functionally annotated the 166 co-expression modules using g:Profiler^38^, which performs functional enrichment analysis on gene sets using predefined gene ontology and pathway annotations. This allowed us to annotate 131 of the 166 modules with at least one significantly enriched gene ontology or pathway (Table S7).

These annotations helped us interpret the function of identified *trans* effects. For example, the *trans*-eQTL signal near *IKZF1* (on chromosome 7) is significantly associated with 27 gene modules. *IKZF1* encodes a transcription factor IKAROS that belongs to the family of zinc finger DNA binding proteins^39^. The *IKZF1* (IKAROS) *trans*-target gene module 159 (M159) is significantly enriched in the “positive regulation of transcription of Notch receptor target” (adjusted *P* = 6.82 x 10^-3^, **Figure 3E**). Reassuringly, it was previously found that IKAROS is a repressor of many Notch targets, and our *trans*-eQTL signal further supports the *trans* regulation of Notch signaling pathway by IKAROS^40^. *IKZF1 trans*-target module 3 (M3) is significantly enriched in the gene ontology term “defense response to virus” (Figure S4, adjusted *P* = 8.7 x 10^-31^) and M35 is significantly enriched in the innate immune system (adjusted *P* = 4.09 x 10^-17^). This data supports the conclusion that the *IKZF1* locus plays a *trans*-regulatory role in immune responses (**Figure 3E**). The *trans*-eQTLs near *NFKBIA*, which encode NF-kappa-B inhibitor subunit A, are significantly associated with module 66 (M66) (adjusted *P* < 1.8 x 10^-7^, adjusted *P* < 9.9 x 10^-2^). Interestingly, we found that M66 is highly enriched in NF-kappa-B signaling pathway (adjusted *P* = 8.35 x 10^-s^, **Figure 3E**), which supports the *trans* regulation of the NF-kappa-B signaling pathway by *NFKBIA*. The complete list of *trans*-eQTLs signals and functional annotations of *trans*-target gene modules can be found in Supplementary Table S4 and Table S7.

### Trans-PCO identifies 965 *trans*-eSNP–module pairs associated with well-defined biological processes

To further demonstrate the utility of trans-PCO, we applied trans-PCO to 50 MSigDB hallmark gene sets, which represent well-defined biological processes^30^, including DNA repair, coagulation, heme metabolism, Notch signaling etc. (Table S15). Each gene set contains between 32 and 200 genes. In DGN, we identified 965 significant *trans*-eSNP–module pairs, corresponding to 41 gene sets and 120 *trans*-loci–module pairs (R2<0.2), at 10% FDR level (Figure S5, Table S3, Table S16).

*Trans*-eQTLs associated with well-defined biological processes facilitate interpretation of the *trans*-eQTL signals. For example, we identified several *trans*-eQTL signals at the *NLRC5* locus (Table S16). The *trans* target gene set is the “interferon alpha response” gene set, suggesting *trans* regulation from *NLRC5* to the interferon signaling pathway. Reassuringly, earlier studies have confirmed that *NLRC5* is a master regulator for MHC class II genes and negatively regulates the interferon signaling pathway^41,42^. The *trans*-eQTL signals also validated our previous interpretations of *trans*-eQTLs associated with co-expression gene modules. For example, in agreement with our analysis of co-expression modules, we found that the *IKZF1* locus is significantly associated with several immune-related biological processes, such as interferon gamma response (Table S16, **Figure 3E**).

**Figure 3.**
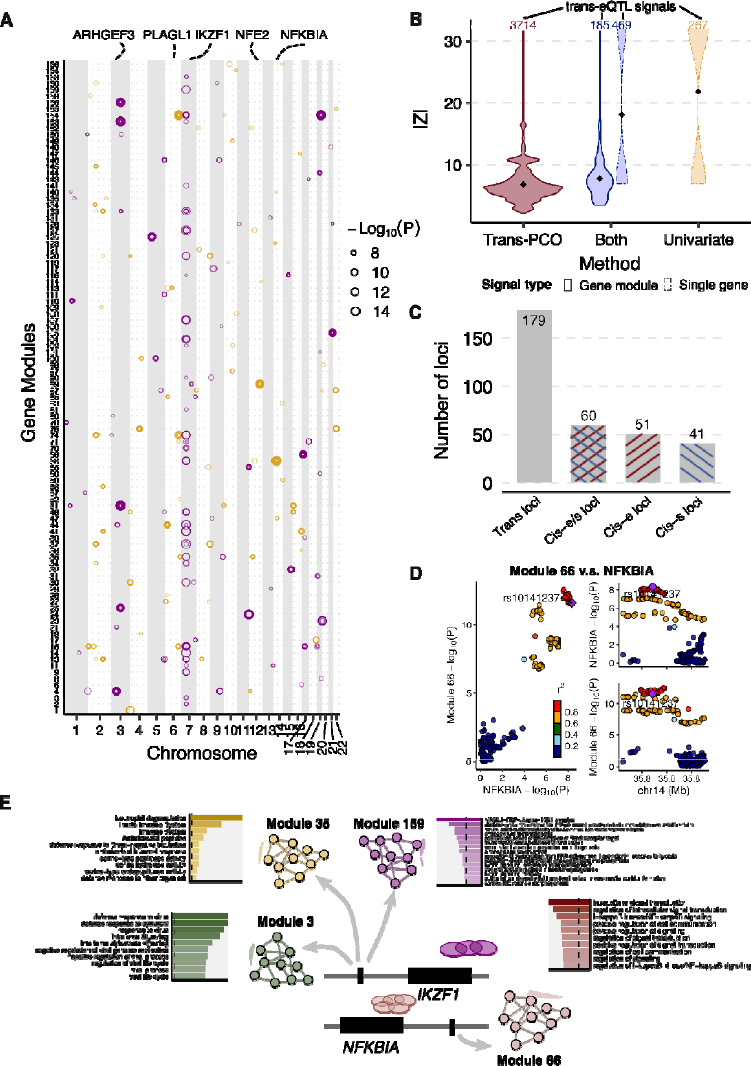
Trans-PCO identifies *trans*-eQTLs associated with co-expression gene modules in DGN. (A) Significant *trans*-eQTL signals associated with 166 co-expression modules in DGN. Chromosomal positions of *trans*-eSNPs are on the x-axis, and gene modules are on the y-axis. Point sizes are -Log_10_(*P*) values of significant *trans*-eQTLs. Purple and orange represent odd and even chromosomes, respectively. (**B)** Comparison of the magnitude of significant *trans*-eQTLs effects detected by trans-PCO and the univariate method. X-axis shows signal categories: trans-PCO specific signals (Trans-PCO), univariate test specific signals (Univariate), and signals identified by both methods (Both). The maximum z-scores of each SNP and each gene in a gene module is used to represent the SNP-module pair. The numbers on top are the number of signals in each category. Line type represents the target type of signals (gene module vs single gene). Y-axis is the absolute value of the z-scores of the signals. (C) Colocalization of *trans*-eQTLs and *cis*-e/sQTLs. The gray bar represents the *trans*-loci used for colocalization analyses. The bar highlighted in blue represents *trans-*loci colocalized with *cis*-sQTLs, red for *cis*-eQTLs, and mixed color for either *cis*-eQTLs or *cis*-sQTLs. (D) Colocalization of *trans*-eQTLs of Module 66 and *cis*-eQTLs of *NFKBIA*. (E) Functional annotations of gene sets facilitate functional interpretation of *trans*-eQTL signals. The *trans*-eQTLs near *NFKBIA* and *IKZF1* are associated with several gene modules. The bar plots show the functional enrichments in modules. The numerical values of enrichments are in Table S7.

### Trans-PCO improves understanding of *trans* regulatory effects of disease-associated loci

To understand *trans* regulatory effects of genetic variants associated with complex traits, we performed colocalization analysis of *trans*-eQTL signals with GWAS loci of 46 complex traits and diseases, including 29 blood traits and 8 other common complex traits (such as height and BMI) from the UK Biobank^37,43^ and 9 autoimmune diseases^34,44–50^ (Methods and Table S8).

We grouped the *trans*-eSNPs into 200kb regions (or *trans*-loci) for colocalization analyses (see Methods). The 3899 *trans*-eQTLs associated with co-expression gene modules were grouped into 179 *trans*-region–module pairs. 42 out of 46 complex traits have at least one GWAS loci colocalized with one of 179 *trans*-region–module pairs. On average across all traits, 8.8% of *trans*-loci colocalize with GWAS loci (**Figure 4A**, Table S9). We observed a higher proportion of colocalization with blood traits (mean proportion 12.0%) than non-blood traits (mean proportion 1.5%). Although we expect some higher proportions of colocalization with blood traits to occur in a whole blood sample, our results may also indicate some residual effects due to cell composition--despite corrections for cell composition using both gene expression PCs and estimated cell-type proportions^27^, such that some *trans*-eQTLs may regulate the abundance of cell proportions and therefore are associated genes that are specifically expressed in certain cell types. Our results are consistent with a recent study by the eQTLGen consortium, which has shown that *trans*-eQTLs in whole blood reflect a combination of cell-type composition and intracellular effects^10^.

**Figure 4.**
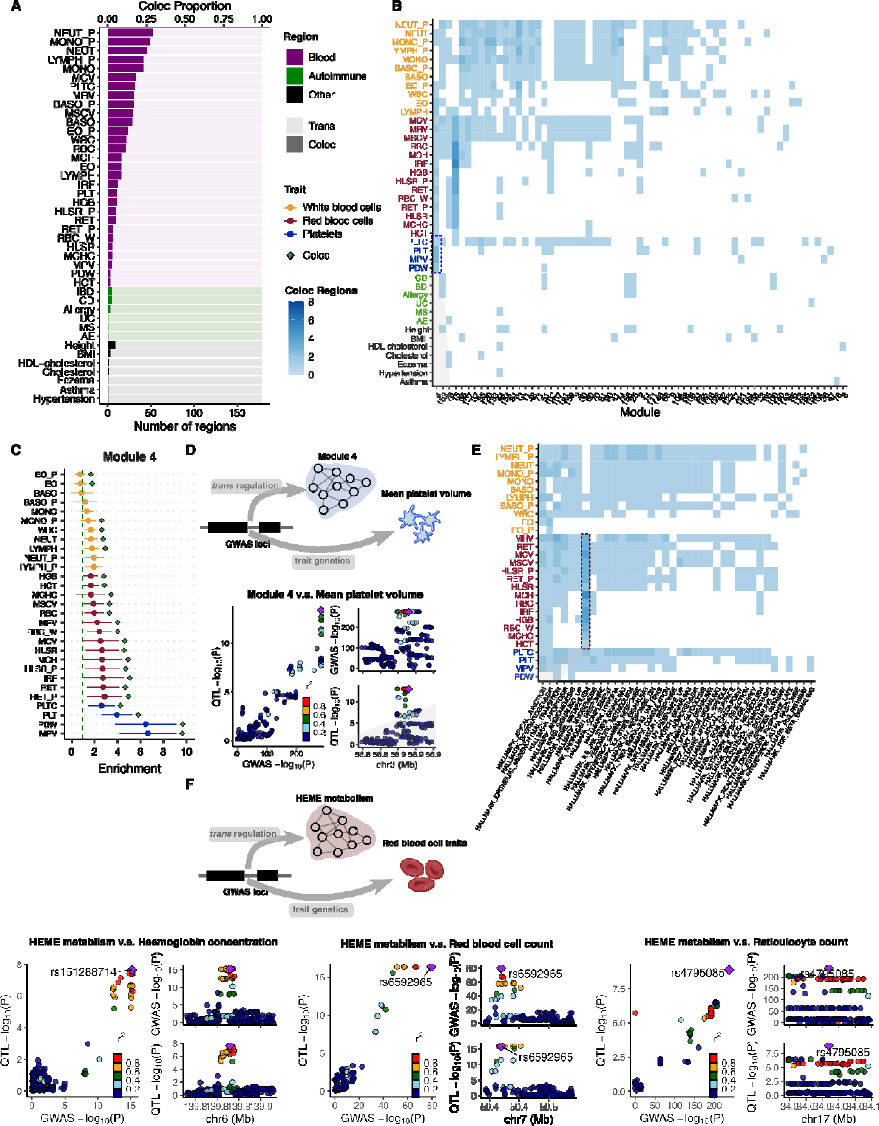
Colocalization of *trans*-eQTLs with GWAS loci of 42 complex traits with at least one colocalization region. **(A) The number of colocalized *trans*-loci associated with co-expression gene modules with GWAS loci.** The traits are first ordered by broad categories: blood traits, autoimmune diseases, and other traits in UKBB. The traits within each category are then ordered by the total number of colocalized regions. (B) Heatmap of the number of colocalized *trans*-loci associated with co-expression gene modules with GWAS loci between each module and trait. The traits are first ordered by broad categories: white blood cells (Orange), red blood cells (Red), platelet cells (Blue), autoimmune diseases (Green) and other traits in UKBB (Black). The traits within each category are then ordered by the number of colocalized gene modules. The blue shades represent the number of colocalized regions. **(C) Heritability enrichment of Module 4 (M4) in blood traits.** Heritability enrichment was estimated by using S-LDSC. Error bars are 95% confidence intervals. (D) Colocalization of mean platelet volume associated locus near *ARHGEF3* and *trans*-eQTL of M4. (E) Heatmap of the number of colocalized *trans*-loci associated with MSigDB hallmark gene sets with GWAS loci across modules and blood traits. The blue shades represent the number of colocalized regions. (F) Colocalization of GWAS loci associated red blood cell traits and *trans*-eQTLs associated with heme metabolism. Six loci associated with red blood cell traits are associated with heme metabolism in *trans*. Numerical results can be found in Table S17. Colocalization plots of the other loci are in Figure S6.

Nevertheless, we found several *trans*-eQTLs that colocalized with GWAS loci, which revealed specific interpretable pathways or functional gene sets (**Figure 4B**, Table S10). For example, *trans*-eQTLs associated with co-expression module 4 (M4) colocalized with 24 out of 29 blood traits (**Figure 4B**). M4 is highly enriched for genes involved in platelet activation (adjusted *P* = 1.12 x 10^-12^, Figure S4, Table S7). One of the colocalized *trans*-eSNPs associated with M4 is in the introns of the *ARHGEF3* gene (**Figure 4D**), which has been shown to play a significant role in platelet size in mice^51^. To further support the interpretation of colocalized signals, we estimated heritability enrichment of M4 in blood traits using stratified LDscore regression^52^ (S-LDSC, **Figure 4C**). We reasoned that an enrichment of trait heritability near genes in a module would strongly support the involvement of a module in the genetic etiology of a trait. Strikingly, we found that M4 is significantly enriched in the heritability of multiple blood traits, and the enrichment was especially strong for platelet traits such as platelet distribution width (enrichment = 6.5 x, *P* = 7.0 x 10^-s^) and mean platelet volume (enrichment = 6.7 x, *P* = 1.2 x 10^-s^, **Figure 4C**, Table S11). Additionally, we evaluated whether M4 genes are significantly enriched in genes associated with platelet traits, identified by transcriptome-side association studies (TWAS). There are 1339 unique genes significantly associated with platelet traits in the UK Biobank^53^. M4 genes are significantly enriched in TWAS genes associated with platelet traits (88 overlap genes, p-value=6.7x10^-10^, Fisher’s exact test), which further supports the role of M4 in platelet traits. Finally, we identified that the *ARHGEF3* locus is significantly associated with the MSigDB coagulation hallmark gene set (Table S16). These findings strengthen the model where genetic variation near *ARHGEF3* impacts the expression levels of multiple genes that are involved in platelet biology and that also harbor nearby genetic variation associated with platelet traits.

We also performed colocalization analysis of *trans*-eQTLs associated MSigDB hall mark gene sets **(Figure 4E**, Table S17). One of the gene sets represents heme metabolism, which is an essential process underlying erythroblast differentiation and red blood cell counts. We found that six *trans*-eQTL loci of heme metabolism significantly colocalized with GWAS loci associated with red blood cell traits, such as hemoglobin concentration, red blood cell count, and reticulocyte count (PP4=0.76-1.00, **Figure 4F**, Figure S6, Table S17). We found that the genes in the gene sets are significantly enriched in TWAS significant genes associated with hemoglobin levels in the UK Biobank (35 overlap genes, p-value=8.1x10^-4^, Fisher’s exact test), which further supports the role of the hallmark gene set in red blood cell traits. Our results provide evidence that these six loci regulate heme metabolism in *trans,* which is an essential process underlying erythroblast differentiation and red blood cell counts.

In another example, we found a *trans*-eQTL near *IKZF1* for M3 that colocalizes with 11 blood traits, seven of which are related to white blood cells (Table S10). As mentioned previously, M3 is significantly enriched for gene ontology terms including “defense response to virus” (adjusted *P* = 8.7 x 10^-31^) and “negative regulation of viral processes” (adjusted *P* = 1.07 x 10^-17^, Table S7). The enrichments are driven by many genes related to interferon (e.g., *IFI6*, *IFI16*, *IRF7*), which are proteins released by host cells in response to the presence of viruses and indicate immune related functions (Table S1, Table S7). Additionally, our heritability analysis of genes in M3 identified enrichments for multiple traits associated with blood cell-type count including neutrophil count (enrichment = 2.3x, *P* = 1.7 x 10^-4^) and white blood cell count (enrichment = 2.1x, *P* = 1.3 x 10^-4^, Figure S7). Our analyses support that the white blood cell associated locus *IKZF1* regulates immune response pathways in *trans*.

Taken together, our functional map of *trans*-eQTLs revealed concrete examples where genetic variants associated with complex traits also influence a biological pathway or a coherent set of genes with similar functions. Thus, *trans*-eQTL of gene sets have the potential to reveal *trans*-regulatory mechanisms underlying complex traits and diseases. The complete list of colocalization signals for each trait can be found in Supplementary Table S10.

### Summary-statistics–based trans-PCO identified 10,167 *trans*-eSNP–module pairs in eQTLGen

We developed summary-statistics–based trans-PCO to increase its applicability to gene expression datasets of large sample sizes, such as eQTLGen^10^ (N=31,684, whole blood). To ensure that summary-statistics–based trans-PCO signals are well-controlled for test statistics inflation and false positives, we added two steps to the original pipeline. First, we carefully select gene sets to minimize the noise when approximating the gene correlation matrices. When only summary statistics are available, the correlation matrix of each gene set is approximated with the correlations of z-scores of the insignificantly associated SNPs of each gene. A low ratio of SNPs to genes (<50) results in a noisy approximation of correlation matrices and test statistics inflation (Figure 6A, Methods, and Supplementary Note). Therefore, we only use gene modules with ratios greater than 50 to test for *trans*-eQTLs, which we show are well-controlled for inflation (**Figure 5A**, Figure S8). Second, we remove genes in the module that are cross-mappable to the test SNP loci (see details in Methods) in the association test to reduce false positives caused by multi-mapping reads.

The eQTLGen study performed the standard univariate *trans*-eQTL mapping on a subset of 10,317 GWAS SNPs and the summary statistics of these *trans*-eQTLs are available. We applied the summary-statistics–based trans-PCO to these summary statistics to identify *trans*-eQTLs associated co-expression gene modules and MSigDB hallmark gene sets.

Of the 166 co-expression gene modules identified in DGN, we used 129 modules with reliable correlation matrix approximations to ensure the *trans*-eQTL signals are well-controlled for inflation (**Figure 5A**, Figure S8, Methods, and Supplementary Note). Similarly, Of the 50 MSigDB hallmark gene sets, we only used 11 gene sets with accurate correlation matrix approximations (Figure S17). In total, there were 4533 genes in the tested co-expression gene modules and hallmark gene sets. For co-expression gene modules, we identified 8116 *trans*-eSNP–gene co-expression module pairs, corresponding to 2161 eQTLGen test SNPs and 122 gene modules (**Figure 5B**, Table S3, Table S12). For hallmark gene sets, we found 2051 significant *trans*-eSNP–hallmark gene set pairs, corresponding to 1018 SNPs and all 11 hallmark gene sets, using Bonferroni correction (Table S3, Table S18). In eQTLGen, we did not perform LD clumping on *trans*-eSNPs, because they were GWAS SNPs associated with different traits and diseases. The univariate method used in eQTLGen^12^ identified 1050 hub SNPs targeting more than 10 genes at 5% FDR, 89% of which are also identified by trans-PCO (**Figure 5C**).

The large sample size in eQTLGen improves the power of *trans*-eQTL detection. Of the 3899 significant *trans*-eSNP–co-expression module pairs in DGN, 38 pairs were also tested in eQTLGen. Reassuringly, we found that all 38 *trans* signals were replicated in eQTLGen (under a replication P-value cutoff of 0.1/38, Table S13) and all association p-values were highly significant (*P* < 10^-12^, Figure S9). In contrast, most of the *trans*-eQTL signals in eQTLGen were not found in DGN. For example, of the 7577 SNP-module pairs analyzed in both datasets, there were 7291 pairs (96%) that were uniquely identified in eQTLGen (which is defined as at least 1 MB away from *trans*-eQTL SNPs in DGN). This is not surprising, because the association p-values are much smaller in the eQTLGen dataset due to the larger sample size (Figure S10). Similarly, eight significant *trans*-eSNP–hallmark gene set pairs in DGN were tested in eQTLGen, and all of them were replicated. We also compared eQTLGen signals by trans-PCO to those identified by ARCHIE in Dutta et al.^24^ (see Supplementary Note and Figure S19).

The nearest genes of eQTLGen *trans*-eQTLs are significantly enriched in DNA binding activity (adjusted *P* = 3.73 x 10^-4^) and transcription factor binding (adjusted *P* = 1.74 x 10^-7^) , as well as immune responses, such as cytokine receptor activity (adjusted *P* = 7.27 x 10^-7^) or MHC class II receptor activity (adjusted *P* = 9.93 x 10^-s^, **Figure 5B**, Table S14). We found that the enrichment of immune responses was driven by *trans*-eQTLs in the HLA region on chromosome 6 (such as *HLA-DRA*, *HLA-DRB1 etc*, Table S12) or near cytokine receptor genes (such as *IL23R, IL1R1, CXCR4 and* genes on the chemokine receptor gene cluster region*: CCR2, CCR3, CCR5 etc*). These *trans*-eQTLs are associated with several autoimmune diseases, such as type 1 diabetes, autoimmune thyroid diseases, cutaneous lupus erythematosus and inflammatory bowel disease (Table S12). The trans-PCO signals help us understand the *trans* regulatory mechanism of these loci. For example, we found that the *trans* target gene modules of the HLA loci are enriched in immune related functions, such as cytokine production (M44), B cell differentiation (M54), IgE binding (M60), TNF signaling pathway (M62), T cell activation (M63 and M87), and cytokine signaling pathway (M62 and M76, **Figure 5D**). The *IL23R* locus is associated with cytokine signaling pathway (M76) in *trans*. The chemokine receptor genes were associated with several gene modules including cytokine production (M44), IgE binding (M60) and T cell activation (M87). These *trans*-eQTL signals support the conclusion that genetic loci associated with autoimmune disease regulate immune related pathways in *trans*.

**Figure 5.**
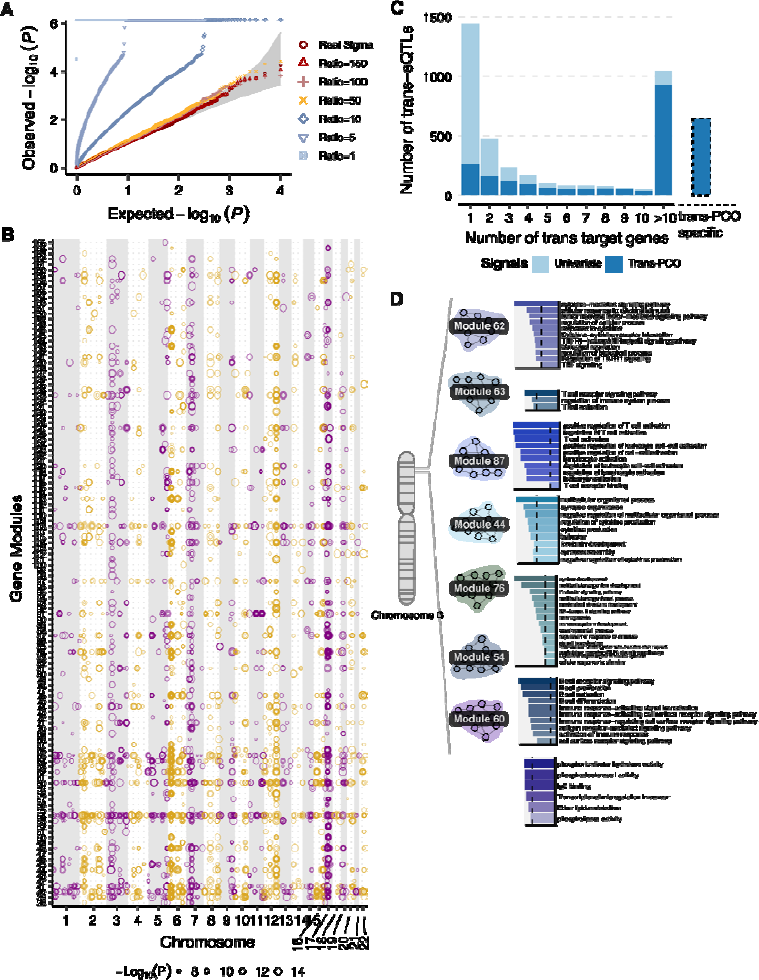
Trans-PCO identifies *trans*-eQTLs associated with co-expression gene modules and MSigDB hallmark gene sets in eQTLGen. **(A) Summary-statistics–based trans-PCO is well controlled for test statistics inflations.** We show gene module 1 (size 625) as an example. SNP to gene ratios used for correlation matrix estimation are in different shapes and colors. Red-yellow shades represent higher ratios (>=50) and blue shades represent lower ratios. Gray area shows 95% CI. Trans-PCO used a minimum ratio of 50. (B) 8199 significant *trans*-eSNP–module pairs associated with co-expression modules in eQTLGen. Chromosomal positions of *trans*-eSNPs are on the x-axis and gene modules are on the y-axis. Point sizes are -Log_10_(*P*) values of significant *trans*-eQTLs. **(C) The majority of hub SNPs targeting more than 10 genes in the original eQTLGen study are identified by trans-PCO.** Light blue bar represents the total number of *trans*-eQTLs in the original eQTLGen study at 5% FDR level. Dark blue bar represents the *trans*-eQTLs also detected by trans-PCO under Bonferroni correction that are associated with co-expression modules or MSigDB gene sets. The bar on the right shows the *trans*-eQTLs detected only by trans-PCO. (D) The HLA locus is associated with several immune related gene modules in *trans*. The bar plots show the functional enrichment of co-expression gene modules.

## Discussion

In summary, we developed a powerful method, trans-PCO, to detect *trans*-eQTLs associated with expression levels of co-expressed or co-regulated genes. The multivariate approach of trans-PCO can detect much smaller *trans* effects (**Figure 3B**, Figure S13) and is substantially more powerful than existing methods. (**Figure 2**). Trans-PCO is also flexible. It can be applied to both RNA-seq data with genotypes or summary statistics, and the user can employ various definitions of gene sets. Applying trans-PCO to both the DGN and the eQTLGen datasets, we identified nearly 15,000 *trans*-eSNP–module pairs associated with co-expression modules and well-defined biological processes. *Trans*-eQTLs with annotated gene modules facilitate our understanding of the *trans*-eQTL signals. These *trans*-eQTLs also improve our understanding of the *trans* regulatory effects of disease associated loci. We highlight multiple examples where our map of *trans* effects helps us identify how trait-associated variants impact gene regulatory networks and pathways. For example, we found six genetic loci associated with red blood cell traits to have significant *trans-*associations with the heme metabolism gene set. It is possible that these genetic loci are “peripheral master regulators” that regulate core processes of red blood cell production.

We thoroughly compared the performance of trans-PCO versus other methods, such as the PC1–based method by Kolberg et al.^23^, ARCHIE by Dutta et al.^24^ and Rovital et al.^22^. Trans-PCO and the PC1–based method are both designed to identify individual *trans*-eQTLs of any gene sets containing multiple genes, and the comparison between them is straightforward in both simulations and real data analyses. However, ARCHIE is different and not directly comparable to the other two methods for several reasons (see more discussions in Supplementary Note). First, ARCHIE captures only trait-specific *trans* regulations. It identifies sets of gene-expressions trans-regulated by sets of known trait-related genetic variants. In addition, ARCHIE tests significance against a competitive null hypothesis, which uses cc-values of all eQTLGen trait-associated variants as empirical null distribution and reflects *trans* regulations not specific to any trait. Therefore, an ARCHIE p-value reflects significance of trait-specific patterns. In contrast, trans-PCO identifies *trans*-eQTLs under the general null hypothesis assuming no *trans* effects. Therefore, trans-PCO can be used to generate comprehensive maps of *trans*-eQTLs in tissues and cell types, in non-trait-specific manner. Using ARCHIE to perform genome-wide scan of *trans*-eQTLs in a non-trait specific manner can be challenging, as non-trait specific p-value is not computable in current implementation of the method and it will be extremely computational challenging (due to the computational intensive resampling procedure and difficulty of manipulating whole-genome LD matrices). Second, trans-PCO and ARCHIE are designed to capture different *trans* regulatory effects. ARCHIE is powerful when multiple disease–associated variants have weak effects on a single gene (for example, multiple GWAS variants converge onto the core genes through *trans* regulation) or multiple disease–associated variants have weak effects on multiple genes (Figure 2 in Dutta et al.^24^), in which multiple genes are not co-regulated by a shared *trans* genetic locus. In contrast, trans-PCO is designed to capture weak *trans* signals of a variant on multiple co-regulated genes, for example, a transcription factor has *trans* effects on multiple target genes. We include detailed comparison of the two methods using both simulations and real data analyses (see Supplementary Notes and Figure S19). Our results support that the two methods are powered at detecting different *trans* signals. Third, ARCHIE identifies components, consisting of multiple trait-associated SNPs and multiple genes, where sets of gene expressions are *trans* regulated by sets of trait-associated variants. Without knowing the exact *trans*-eQTL SNP driving the *trans* regulation, it is hard to further study *trans* regulatory mechanisms of the *trans*-eQTLs, for example, whether the *trans-*eQTL is also a *cis-*eQTL, or which gene is the *trans* regulator. Fourth, ARCHIE takes all genes as input and infers gene sets that are *trans*-regulated by disease-associated variants, whereas trans-PCO is flexible to be applied to any user-defined gene set of interest to identify *trans*-eQTLs. The genes in the ARCHIE components are likely “core” genes for a specific disease and can be used to find key biological processes for the disease. Trans-PCO could also be used to identify disease relevant genes and processes through follow up analyses, such as colocalization analyses. In summary, trans-PCO and ARCHIE have different goals and are designed for detecting different types of *trans* signals. Yet, we thoroughly compared ARCHIE and trans-PCO in both simulations and real data analyses (Supplementary Note), (1) in simulations, we evaluated whether ARCHIE can identify regular *trans*-eQTLs detectable by other methods (trans-PCO, PC1–based and univariate method), (2) in real data analyses (eQTLGen summary statistics), we evaluated whether trans-PCO can identify *trans*-signals identified by ARCHIE. We believe these comparisons will provide insights on when and how these methods should best be used. In addition, Rovital et al.^22^ used independent component analyses to identify components representing co-expression patterns from the expression of all genes, and identified *trans*-eQTLs that have enriched *trans* associations with the components. However, we demonstrated through simulations that the Rotival et al. method has minimal power to identify weak *trans* effects (Supplementary Note, Figure S20).

A limitation of multivariate association tests, including trans-PCO, is that they do not explicitly identify which genes in the gene sets are significantly associated with the test SNP. While functional annotations of gene sets facilitate our understanding of the *trans*-eQTL signals, it is possible that the genes driving *trans*-associations are different from the genes driving functional enrichment of the gene sets. Therefore, the biological interpretation of *trans*-eQTL signals should be supported with other evidence before it is considered definitive. However, there are exploratory analyses that can help prioritize genes in the network that are key drivers of the underlying signal. For example, by examining the univariate association p-values between the *trans*-eQTL SNP and each gene in the network, the user can prioritize genes with the most significant p-values as likely *trans* targets. Furthermore, the users can also use the rr1 statistics on the univariate p-values to estimate the proportion of genes that have true *trans* effects in the network. While the exact molecular mechanism requires further validation, the large number of *trans*-eQTLs identified by trans-PCO in our study opens up new opportunities to understand complex traits-associated loci and underlying mechanisms.

*Trans*-eQTLs identified in bulk tissues can be a combination of cell composition *trans*-eQTLs, which are driven by cell type proportions, and intracellular *trans*-eQTLs, which capture *trans* regulatory effects in a single cell type. To get higher proportions of intracellular eQTLs, the common approach is to correct for cell type proportions in association tests. For example, the eQTLGen study^12^ corrected for cell proportion effect by using gene expression PCs. They validated some trans-eQTLs using single-cell RNA sequencing data, indicating that these *trans*-eQTLs were intracellular *trans*-eQTLs. In our analysis of DGN dataset, we included the estimated cell proportions as covariates, in addition to gene expression PCs. This strategy might have given rise to higher proportions of intracellular *trans*-eQTLs. Co-expression gene modules could also capture cell proportion effects. In our study, we removed cell proportions from gene expression levels before clustering genes into co-expression modules. While this can correct for cell proportion effects in the co-expression modules to some extent, we note that it does not guarantee their complete removal.

Many studies, including ours, seek to avoid cell composition effects. However, by closely examining *trans*-eQTLs discovered in our study, we think cell composition *trans*-eQTLs can be biologically interesting too. For example, the *IKZF1* locus is significantly associated with several gene modules enriched with viral defense and other immune related functions in *trans*. The locus is also significantly associated with white blood cell proportions. Given the general function of white blood cells in fighting infections, these observations raise the possibility that the *trans*-eQTLs near *IKZF1* regulate antiviral activity by affecting white blood cell-type proportion. Supporting this hypothesis, we found earlier that genetic variants near *IKZF1* are also associated with expression levels of genes in M159, which are enriched in genes involved in the Notch signaling pathway. The Notch signaling pathway plays a central role in cell proliferation, cell fate, and cell differentiation^54^; thus, our analyses reveal a plausible mode of action whereby genetic variants near *IKZF1* impact multiple immune-related functions by influencing white blood cell-type proportions. In future studies, it could be interesting to specifically identify cell proportion effects and understand their role in complex traits.

Identifying the network effects of genetic variants not only shed light on molecular mechanisms of complex associated loci, it can also have important translational applications, for example, in drug discovery and development. First, genes that are associated with disease relevant pathways can serve as evidence for therapeutic targets of the disease. In a preliminary analysis, we examined whether allergy drug targets are more likely to be associated with immune-related gene sets. Among a total of 142 gene sets (129 co-expression gene modules and 11 hallmark gene sets) used for *trans*-eQTL identification in eQTLGen, 19 were defined as immune-related. We used 55 launched allergy drug target genes from The Broad Institute Drug Repurposing Hub (https://repo-hub.broadinstitute.org/repurposing), 5 of which are near allergy associated loci in eQTLGen. Interestingly, we found all 5 targets to be associated with immune-related gene sets (Table S19). Detailed analyses can be found in Supplementary Note. While the enrichment is not statistically significant (P=0.12, Fisher’s exact test; Table S20), it is likely due to the small number of drug targets included in our analyses. Additionally, we observed that the *trans* gene modules of drug targets converge to gene sets whose functions are highly relevant to allergy. For example, three drug targets (*IL3*, *UGT3A1* and *SLC37A4*) are associated with gene sets enriched for the B cell signaling pathway. More comprehensive analyses are beyond the scope of this study, yet our preliminary analyses have demonstrated that one can consider genes that have strong *trans* associations with disease-relevant pathways to identify drug targets for the specific disease, especially those with known disease-relevant pathways. Second, network effects of disease variants can be used for repurposing existing drug compounds to new diseases. Drug repurposing can substantially reduce cost and time to develop new treatments. If the gene expression profiles of an existing drug is enriched for genes in the *trans*-network of another disease’s associated loci, it can serve as an evidence for repurposing. Additionally, knowing the network effects of a gene can also help evaluate the safety of a potential drug target. Therapeutic perturbation of a drug target can affect expressions of many downstream genes. While some of them are in the desired disease pathways, others are in pathways associated with other phenotypes, inducing unwanted side-effects. We believe comprehensive catalogs of *trans*-networks effects in human cell types and tissues will serve as important resources for interpretation of *trans* regulatory effects of disease associated loci as well as translation applications. Therefore, we made all the trans-PCO *trans*-eQTL signals, with functional annotation of the gene sets, publicly available, downloadable and browsable in www.networks-liulab.org/transPCO.

## Methods

### Trans-PCO pipeline

We test if a genetic variant is associated with genes in a module through *trans* regulations using the multivariate model as follows,

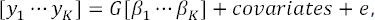

where *G* is the dosage of a reference allele representing the genotype of a SNP, *β_k_* is the effect of the SNP on *k*-th gene in the module with *K* genes, and *y_k_* is the expression level of the *k*-th gene. To test if a SNP of interest is significantly associated with the module, we test the null hypothesis,

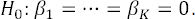

We use a PC-based omnibus test (PCO)^26^, which is a powerful and robust PC-based approach aiming at testing genetic association with multiple genes with no prior knowledge of the true effects.

Specifically, PCO combines multiple single PC-based tests in linear and non-linear ways, corresponding to a range of causal relationships between the genetic variant and genes, to achieve higher power and better robustness. A single PC-based test (most commonly the first primary *PC*_1_) is,

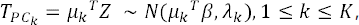

where *z* is a *K* × 1 vector of univariate summary statistic *z*-scores of the SNP for *K* genes in a module, *µ_k_* is the *k*-th eigenvector of the covariance matrix 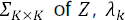 is the corresponding eigenvalue, and *β* represents the true causal effect. PCO combines six PC-based tests, including,

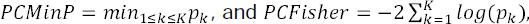

where *p_k_* is the p-value of *T_PC_k__*. These two tests take the best p-value of single PC-based tests and combine multiple PC p-values as the test statistic. Other tests include,

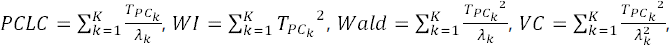

which are linear and quadratic combinations of each single PC-based test weighted by eigenvalues. The six tests achieve best power in specific genetic settings with different true causal effects^26^. PCO takes the best p-value of the PC-based tests as the final test statistic,

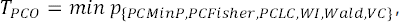

to achieve robustness under unknown genetic architectures while maintaining a high power. The p-value of PCO test statistics can be computed by performing an inverse-normal transformation of the test statistics,

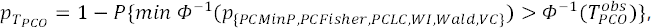

where Φ^-1^ denotes the inverse standard normal cumulative distribution function. The p-value can be efficiently computed using a multivariate normal distribution as described in Liu et al.^26^.

To prevent cis-regulatory effects from driving the identified *trans* associations between a SNP and module, we removed genes in the module that are on the same chromosome as the tested variant. In addition, to avoid false positive signals in *trans* associations due to alignment errors, we discarded RNA-seq reads that are mapped to multiple locations or poorly mapped genomic regions (mappability score <1)^27,28^ before quantifying gene expression levels.

### Simulation

To evaluate the power of trans-PCO, we performed a series of simulations with various parameter settings corresponding to different genetic architectures. We applied trans-PCO to the simulated datasets and assessed the false positive rate and statistical power. We also ran two additional statistical tests, a univariate test (“MinP”) and a primary PC-based test (“PC1”), to compare their performances with trans-PCO and to obtain more in-depth insights on trans-PCO,

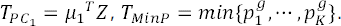

The PC1 based test takes only the first PC as the proxy of a gene module and uses it as the response variable to test for genetic variants with significant associations. We also compared trans-PCO with a non-PC based statistical test, MinP, which takes the minimum p-value across genes in the module as the test statistics.

To implement trans-PCO, PC1, and MinP tests, there are two main pieces of information that are required as input, i.e. correlation matrix of the gene module and summary statistics (z-scores) of SNPs with genes in the module. We used a gene module from the RNA-seq data (see Genotype and RNA-seq QC) consisting of 101 genes (*K* = 101) and the corresponding correlation matrix to make the settings more realistic. We sampled z-scores of 10^7^ SNPs from the null distribution,

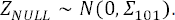

We tested the associations between each SNP and the gene module using the above three tests and evaluated the p-values against the uniform distribution to validate if the statistical tests are well calibrated.

We also simulated 10k z-scores of SNPs from the alternative distribution,

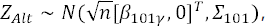

where *n* is the sample size, *β* is a 101y-long vector representing the causal effect of a SNP on 101 genes, and *γ* is the proportion of true target genes in the module. Each component of β follows 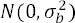), where 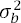 is the genetic variance. By default, we set the sample size *n* to be 500, 30% genes (30) in the module are true *trans* target genes, and 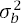 to be 0.001. To evaluate how three tests perform across different genetic architectures, we simulated multiple scenarios across varying sample sizes, target gene proportions, and genetic variances. Specifically, we looked at the cases where sample size is 200, 400, 600, and 800, causal genes proportion is 1%, 5%, 10%, 30%, and 50%, and genetic variance is 0.002, 0.003, 0.004, 0.005, and 0.006. We simulated 10k SNPs and performed 1000 simulations. To control the false discovery rate, we corrected the p-values for multiple testing based on the simulated empirical null distribution of p-values, to keep it consistent with the method used in the RNA-seq dataset (see Supplementary Note). An association is significant if its adjusted p-value is lower than 0.1. The power is calculated as the proportion of SNPs that were identified to be significant among 10k SNPs.

### Genotype and RNA-seq QC

We analyzed an RNA-seq dataset from whole blood^27^. We performed a series of QC on individuals, genotypes, RNA-seq reads, and genes before quantifying gene expression profiles. Specifically, we referred to the procedures in Liu et al.^28^. For individual-level QC, we removed related individuals from 922 samples with RNA-seq reads available and kept 913 individuals in total for further analysis. For genotype-level QC, we used SNPs with genotyping rate >99%, minor allele frequency >5%, and Hardy-Weinberg equilibrium < 10^-6^.

RNA-seq reads can be falsely aligned to genomic regions with high sequence similarity. The misalignment onto these regions can lead to false positive signals in *trans*-eQTL analysis and spurious correlations in gene co-expression networks^8^. To help address this problem, we performed RNA-seq read-level QC to remove the reads with alignment issues. To be more specific, we filtered out the reads that were mapped to multiple genomic regions and reads with >2 mismatches. We also removed the reads aligned to regions with low mappability.

On the gene level, we quantified the gene expression levels as Transcript Per Million (TPM). We first normalized the expression levels across samples to the normal distribution by quantile normalization, and then we normalized the expression levels across genes. We also filtered out genes that are not protein-coding, lincRNA genes, or genes on sex chromosomes. As a result, there are 12,132 genes left for follow up analysis. Finally, to control for potential confounding factors and capture the co-expressed gene modules only driven by genetic effects, we regressed out covariates from the expression profiles. We used biological and technical covariates, including genotype PCs, expression PCs, and blood cell type proportions^27,28^.

### Identification of the gene co-expression network

We are interested in jointly testing co-regulated genes in a multivariate association test. To this end, we first used WGCNA^29^ to construct a gene co-expression network, where genes are connected through correlations among their residualized expression levels. WGCNA uses hierarchical clustering to cut the network into separate gene modules with highly correlated expression levels. We used the default parameter settings, except that we specified the minimum module size parameter (‘minModuleSize’) to 10 to obtain small gene modules.

### Colocalization of *trans*-eQTLs and GWAS loci

To explore the role of *trans*-eQTLs in understanding complex traits and diseases, we performed colocalization between *trans*-eQTLs of a gene module and GWAS loci of 46 complex traits and diseases. Specifically, we used GWAS summary statistics of 29 blood-related traits^37^ and 8 other traits from UK Biobank, provided by Neale Lab (http://www.nealelab.is/uk-biobank/), and 9 autoimmune diseases collected in Mu et al.^34^ (Table S8).

To define a region to perform colocalization, we first selected the *trans*-eQTL with the most significant p-value and expanded a 200kb flanking genomic region centered at the lead SNP as a region to perform colocalization analysis. We then moved on to the next most significant SNP and expanded a 200kb flanking region. We stopped searching for lead SNPs when all *trans*-eQTLs were included. This resulted in 255 *trans* region-module pairs. As two adjacent regions could correspond to the same colocalization signal, we marked adjacent regions as a region group if their lead SNPs were within 200kb, which generated 179 *trans*-region–module pairs in total. We ran colocalization analysis between each 200kb *trans* region and GWAS loci of 46 complex traits using the R package coloc^33^, assuming there is at most one causal variant for each region. We used the default priors and 0.75 as the PP4 cutoff for significant colocalizations. We defined a merged region group as being colocalized with a trait if any of its 200kb sub-regions has significant colocalization with the trait. We visualized the colocalized regions using LocusCompareR^55^.

### Colocalization of *trans*-eQTLs and *cis*-e/sQTLs

We performed colocalization analysis between *trans*-eQTLs and *cis*-eQTLs (*cis*-sQTLs) of genes near the *trans*-eQTLs. We used the same 179 *trans*-region–module pairs defined in the colocalization analysis of GWAS loci. For a *trans* loci, we searched for the genes within 500 kb around the lead *trans*-eQTLs of the loci, and used these genes to perform colocalization. We used summary statistics of *cis*-eQTLs and *cis*-sQTLs in the DGN dataset from Mu et al.^34^. We ran coloc^33^ with default priors and 0.75 as PP4 cutoff.

### Trait heritability enrichment in gene modules

To investigate whether a gene module is enriched for trait heritability, we applied stratified LD score regression^52^ (S-LDSC) to 166 co-expression gene modules and 46 complex traits and diseases. Specifically, for each module we defined the annotation set as the SNPs within genomic regions of genes in the module and also a 500 base-pair window around the genes. We also included 97 annotations from the baseline model. Partitioned heritability enrichment was calculated as the proportion of trait heritability contributed by SNPs in the module annotation over the proportion of SNPs in that annotation.

### Summary-statistics–based trans-PCO applied to eQTLGen

The eQTLGen Consortium^10^ has conducted the largest *cis*- and *trans*-eQTLs association analyses in blood to date. Specifically, 31,684 samples were tested for over 11 million SNPs across 37 cohorts. The summary statistics of *trans*-eQTLs are available for 10,317 trait-associated SNPs on 19,942 genes.

We applied our pipeline trans-PCO to eQTLGen summary statistics, using the same 166 co-expression gene modules defined in DGN dataset. We searched for *trans*-eQTLs among 10,317 SNPs.

The eQTLGen summary statistics are marginal z-scores meta-weighted across multiple cohorts. Most z-scores are from studies where the RNA-seq reads with mappability issues were not filtered out before quantifying gene expression profiles. Therefore, directly applying trans-PCO to the summary statistics can lead to false positive signals, which are driven by the cross-mappability between the genes in the module and the *cis*-gene of the test SNP. In order to reduce false positive *trans* signals, we removed from the gene module genes that are cross-mappable to the *cis*-gene (within 100kb) of the test SNP, which is a common practice used in previous studies^8,27,56^. We further removed genes on the same chromosome as the test SNP to prevent the detected *trans* effects from being dominated by *cis* regulations.

The gene expression profiles are not available in eQTLGen. Therefore, to estimate the gene correlation *Σ* of a module, we searched among eQTLGen SNPs for SNPs insignificantly associated with the module (null SNPs) (see Supplementary Note for details). We observed that there are less null SNPs that can be found for large modules. And simulations show that the low ratio of the number of null SNPs used for *Σ* estimation to the module size leads to false positive signals (Supplementary Note). Therefore, we removed 37 gene modules with ratios lower than 50. Finally, we performed trans-PCO on the remaining 129 gene modules.

## Data availability

All *trans*-eQTL signals, with functional annotation of the gene sets, can be browsed and downloaded at www.networks-liulab.org/transPCO. The genotype and gene expression data of DGN were downloaded by application through the NIMH Center for Collaborative Genomic Studies on Mental Disorders, under the “Depression Genes and Networks study (D. Levinson, PI)”. The eQTLGen summary statistics are publicly available at https://www.eqtlgen.org/. The MSigDB hallmark gene sets are publicly available at http://www.gsea-msigdb.org/gsea/msigdb/human/genesets.jsp?collection=H. GWAS summary statistics of traits in the UK Biobank are available at Neale Lab (http://www.nealelab.is/uk-biobank/).

## Code availability

The trans-PCO pipeline and code to reproduce analyses is available at https://github.com/liliw-w/Trans. This work also uses TensorQTL (https://github.com/broadinstitute/tensorqtl) to perform QTL mapping between genotypes and single genes, R package coloc (https://github.com/chr1swallace/coloc) to perform colocalization analyses between trans-eQTLs and cis-eQTLs, cis-sQTLs and GWAS loci, and S-LDSC software (https://github.com/bulik/ldsc) to estimate trait heritability enrichment in gene modules.

## Declaration of interests

The authors declare no competing interests.

## Acknowledgements

We thank Y. Li, A. Dahl, Y. Gilad and Z. Mu for useful scientific discussions. We thank N. Gonzales, C. Jones and S. Sumner for editing the manuscript. This work was funded through the NIGMS Maximizing Investigators’ Research Award (R35GM138084).

## Notes

### Competing Interest Statement

The authors have declared no competing interest.

### Summary of Updates

Add more comparison between our method and other methods in Results, Discussion, and Supplementary Notes.

