## Supplementary Note for "*Trans*-eQTL mapping in gene sets identifies network effects of genetic variants"

#### Trans-PCO on individual level RNA-seq data

##### Estimating correlation matrix

To implement trans-PCO on a pair of SNP and gene module, we need two pieces of information, (i) z-score vector of the SNP on each individual genes in the module, and (ii) estimated $\Sigma$ of the module.

In the analysis of DGN dataset[1], we calculated the summary statistic z-scores using TensorQTL[2], which performs ultrafast trans-QTL mapping. We included 74 biological and technical factors as covariates, including 5 genotype PCs, 10 expression PCs, and the estimated blood cell type proportions etc[1,3]. In order to search along the whole genome for trans-eQTLs of gene modules, we calculated genome-wide z-scores for every gene included in all modules (over 10k genes).

There are two ways to estimate $\Sigma$ of a gene module[4]. The first way is to use correlation matrix of the residualized gene expression levels,

$$\Sigma = cor(Y|covariates),$$

where $Y|covariates$ is the residual gene expression levels after regressing out covariates. We used this estimation when gene expression profiles are available, as in the case of DGN.

The other way to estimate $\Sigma$ is to use the covariance matrix of insignificant z-scores (null z-scores) across genes in the module[4],

$$\Sigma = cor(Z),$$

where $Z$ is the z-score matrix of null independent SNPs and genes in the module. We used this estimation when only summary statistics are available, as in the analysis of eQTLGen. More specifically, we collected a large set of independent null SNPs, took the z-scores of SNPs across module genes, and calculated the sample covariance matrix using z-scores over the independent null SNPs. See more details in "Summary statistics based trans-PCO".

##### Multiple testing correction

In the case of analyzing DGN dataset, we tested the genome-wide associations of 166 co-expression gene modules. To correct for multiple testing, we used the empirical null distribution. Specifically, we first randomly permuted sample labels and obtained the null summary statistics across SNPs and genes. Then, we calculated the associations for each pair of SNPs and modules using the null z-scores, and used the p-values as the empirical null distribution. Finally, we corrected each observed p-value by counting how many null tests fall below the observed p-value. We did ten permutations and used the average corrected p-values to claim significance (FDR<0.1). By using the empirical null distribution to correct p-values, we are able to control the false positive rate and preserve the LD structure among SNPs as well as the correlation structure within modules.

#### PCs included in trans-PCO

Trans-PCO tests multiple genes jointly by combining multiple PC's of the genes. However, it is not always best to use all PC's, as suggested by Liu et al.[4], due to the tradeoff between statistical power and numerical stability. Particularly, for a large module with highly correlated genes, the correlation matrix $\Sigma$ would be close to being ill-conditioned. Therefore, the eigenvalues of the last few PC's would be very small, which can lead to inflated test statistics. For example, VC test (one of the six tests PCO constructs its test statistics on),

$$VC = \sum_{k=1}^{K} \frac{{T_{PC_k}}^2}{\lambda_k^2},$$

is weighted by the inverse of eigenvalues, where $K$ is the module size, $\lambda_k$ is k-th eigenvalue, and $T_{PC_k}$ is the k-th PC test statistic. The VC test can be numerically unstable if it includes the last few PC's that have very small eigenvalues.

As a matter fact, we observed a few modules with very small eigenvalues (Figure S11). To investigate how including PC's with small eigenvalues affects the association tests, we did simulations where we performed tests incorporating different numbers of PC's. We found that by including PC's with extremely small eigenvalues, the p-values of null tests were inflated. Therefore, instead of combining all PC's, we discarded the last few PC's with extremely small eigenvalues and used only the top PC's with eigenvalues larger than 0.01.

### Summary statistics based trans-PCO

In the case where only summary statistics are available, in order to perform multivariate association test between a SNP and a gene module, we approximated $\Sigma$ of the module by calculating the sample covariance matrix of z-scores over a large set of independent null SNPs[4] (see previous section). More specifically, we selected independent SNPs that are insignificantly associated (*P* < 1e-4, Figure S12) with all genes in the module, collected the z-scores of genes over these independent null SNPs, and calculated the sample correlation matrix.

We applied trans-PCO to eQTLGen summary statistics[5]. Specifically, we grouped eQTLGen genes into 166 co-expression modules as defined using DGN dataset. There are only 10,317 trait associated SNPs analyzed in eQTLGen that have full summary statistics for all genes available. Therefore, we searched for independent null SNPs of modules among the limited set of SNPs. One issue is that less SNPs were found to have insignificant associations with all genes in larger modules, which means a low ratio of independent null SNPs over module size for these modules (Figure S8).

#### Simulations to evaluate and eliminate signal inflations

We wanted to look into if a low ratio of independent null SNPs over module size can lead to noisy estimation of correlation matrix and inflated signals for larger modules. Therefore, we performed simulations to evaluate the p-values distribution of null tests given various $\Sigma$ estimations using a range of ratios.

We chose a gene module of size $K$ from 166 DGN co-expression modules and the corresponding $\Sigma_K$ estimated by DGN gene expression profiles. We first simulated 10k null SNPs with insignificant associations from $Z_K \sim N(0, \Sigma_K)$ as the test set. We then generated z-score matrix $Z_{m\times K}$ of $m$ null SNPs from $\Sigma_K$ ($Z_{m\times K} \sim N(0, \Sigma_K)$) as a training set to be used for $\Sigma_K$ estimation. To look at the how using various ratios of independent null SNPs over module size ($m/K$) affects signal identification, we estimate a series of $\widehat{\Sigma_K}$ using the sample correlation of $Z_{m\times K}$ under various number of null SNPs ($m$). Lastly, we tested the 10k null SNPs by applying trans-PCO using the estimated $\widehat{\Sigma_K}$ by various $m/K$ ratios. To look at how $\widehat{\Sigma_K}$ affects trans-PCO p-values, we plotted QQ-plot of p-values of all $m/K$ ratios. We performed simulations for modules of various sizes, including module 1-11, 15, 20, 30, 40, 50, 60, 70, 90, 100, 150, 166. We used various $m/K$ ratios, including 1, 5, 10, 50, 100, 150 (Figure S8).

We observed that low $m/K$ ratio can result in inflated null signals, especially for ratios under 50. Therefore, in order to control signal inflations and avoid false positive signals, we removed those large modules with low $m/K$ ratios under 50 (Figure S8) from the following trans-eQTLs detection. As a result, we removed 37 co-expression gene modules and performed trans-PCO on the remaining 129 modules.

#### Other modifications to summary statistics based trans-PCO

We made several other modifications to trans-PCO to make it feasible when only summary statistics are available. We used a more conservative multiple testing correction method, Bonferroni correction, to correct for multiple testing. It is not possible to use the permutation based correction as in analyzing DGN dataset, because no individual level data is available. Therefore, we corrected p-values by multiplying 10,317 (the number of tested SNPs) and 129 (the number of tested gene modules).

We also removed genes from the module that are located on the same chromosome as the SNP, in order to avoid *cis* effects. Additionally, we removed genes cross-mappable with any *cis* genes within 100kb of the tested SNP. This is to reduce false positive trans-eQTLs due to possible sequence errors when calculating summary statistics from RNA-seq reads without carefully filtering out problematic reads.

As a summary, we made a few modifications to summary statistics based trans-PCO, to ensure the trans signals are well controlled for test statistics inflation and false positives. First, we estimated the correlation matrix of modules using a large number of independent null SNPs (*P* < 1e-4). Second, we considered only 129 modules that have accurate estimated correlations ($m/K > 50$) for signal detections. Third, we tested the associations between SNPs and genes in the module that are on different chromosomes as the SNPs. Fourth, we removed genes from the module that are cross-mappable with any cis genes of the SNP (<100kb).

#### Apply the primary PC method to DGN dataset

To compare the primary PC method[7] that includes only the first PC in the test and trans-PCO that combines multiple PC's, we also applied the primary PC approach to detect trans-eQTLs of the RNA-seq dataset (DGN). Specifically, the test statistic we used to test a pair of SNP and gene module is,

$$T_{PC_1} = {\mu_1}^T Z,$$

where $Z$ is the z-score vector of the SNP over the gene module, $\mu_1$ is the eigenvector of the first PC. The idea is to utilize the first PC of the module as a one-dimensional proxy phenotype to represent the module and then test its association with SNPs.

The procedure of applying the primary PC method to DGN is similar to that of applying trans-PCO. Specifically, we started with the processed gene expression profiles by removing RNA-seq reads that were poorly mapped or cross-mapped to multiple genomic regions (see Methods). We then regressed out the biological and technical covariates as in trans-PCO. We obtained the first PC from the residual gene expression levels to calculate the test statistic. We also constructed the co-expression gene network from the residualized expression levels, and tested the same set of 166 co-expression gene modules defined as in DGN dataset. We also removed genes from the tested module that are located on the same chromosome as the tested SNP, to avoid the confusion of *cis* effects. We then performed a genome-wide scan of trans-eQTLs of 166 modules using the primary PC method. We corrected for multiple testing based on the empirical null distribution of p-values obtained by permuting sample labels. We did ten permutations. An association is claimed to be significant if the average corrected p-value is under 0.1.

#### Compare trans-PCO and ARCHIE

We compared trans-PCO to the ARCHIE method proposed in Dutta et al.[8]. ARCHIE is a summary statistic-based method with the goal of identifying sets of gene expressions *trans-regulated* by sets of known trait-related SNPs. The key output of ARCHIE is ARCHIE components, which are a set of selected genes and a set of trait-relevant SNPs whose linear combinations have a high canonical correlation (cc-value) due to trait-specific *trans* regulations. Specifically, Dutta et al.[8] analyzed genetic variants associated with 29 traits using eQTLGen summary statistics. The authors made ARCHIE components of three traits (Figure S19) publicly available, including prostate cancer, schizophrenia, and ulcerative colitis.

ARCHIE and trans-PCO are designed with different goals and usages. First, *trans* regulations captured by ARCHIE components reflect only trait-specific associations. ARCHIE uses only variants specific to a specific trait as input and finds *trans* regulations by these variants. Additionally, ARCHIE tests significance against a competitive null hypothesis, which characterizes all trait-associated variants and reflects general *trans* regulations not specific to any trait, as trait-associated variants are expected to be enriched for *trans*-eQTLs[8]. Therefore, a p-value under the competitive null hypothesis reflects the significance of trait-specific patterns. In contrast, trans-PCO identifies all *trans*-eQTLs of given tissues and cell types under the general null hypothesis assuming no *trans* effects.

Second, trans-PCO and ARCHIE are designed to capture different *trans-regulatory* effects. As shown in ARCHIE simulations[8], it is powerful in the case where one gene has multiple weak *trans* effects or more complicated *trans* effects between multiple SNPs and multiple genes. In fact, they observed that a majority of novel genes they found have multiple weak *trans* associations with variants selected in ARCHIE components. In contrast, trans-PCO is powerful in detecting *trans* signals where a SNP has weak *trans* effects on multiple gene expressions as shown in our simulations.

Third, another difference lies in the way gene sets (modules) are defined by two methods for evaluating associations. Specifically, ARCHIE takes all genes as input and infers gene sets as gene components, whereas trans-PCO is flexible to be applied to any user-defined gene set (module) of interest, such as biological pathways or processes.

Lastly, ARCHIE identifies components, consisting of multiple trait-associated SNPs and multiple genes. The interpretation of an ARCHIE component is that sets of gene expressions are *trans* regulated by sets of trait-associated variants, without knowing which exact variant drives the association. ARCHIE components make it useful for identifying genes involved in traits through *trans* regulation, but not to identify *trans*-eQTL SNPs. In contrast, trans-PCO identifies associations between a single variant and multiple genes, which makes it easier to interpret *trans* signals (e.g., whether a *trans*-eQTL is a *cis*-eQTL or *cis*-sQTL, or whether the nearest gene to the top *trans*-eQTL is a transcription factor). Although ARCHIE can also be applied to single-variant cases, the power is extremely low (Figure S19D).

The differences in goals and usages of ARCHIE and trans-PCO make them not directly comparable. However, to give readers and users insights on when and how each method should be used, we compared their performance using three strategies: (1) in simulations, we evaluated how well ARCHIE can detect regular *trans*-eQTL signals between a single SNP and a gene set, and (2) in eQTLGen, we evaluated how well trans-PCO can replicate signals detected by ARCHIE, and (3) we compared *trans* signals reported by trans-PCO and ARCHIE in the eQTLGen datasets. The details of the three comparisons are as below.

#### Comparison of performance in simulations

To evaluate how well ARCHIE can detect regular *trans*-eQTL signals from a single SNP, we applied ARCHIE to our simulation settings as described in Methods. To recapitulate, we used a real co-expression gene module consisting of 101 genes from DGN dataset. We simulated z-scores from a normal distribution using the correlation matrix of the gene module, sample size 500, causal gene proportion 30% with 30 genes being true target genes, and genetic variance 0.001 as parameters. Trans-PCO has a power of 36% under this setting.

To run ARCHIE, three main inputs are needed[8], including (1) $\Sigma_{GG}$, column-correlations of genetic variants, (2) $\Sigma_{EE}$, column-correlations of gene expressions, and (3) $\Sigma_{GE}$, cross-covariance matrix between variants and gene expressions. In our simulation settings, we tested one variant at a time. Therefore, $\Sigma_{GG} = 1$. We set $\Sigma_{EE} = \Sigma_{101}$, and we approximate $\Sigma_{GE}$ with $\frac{Z_{101}}{\sqrt{N}}$. We used various genetic variances ranging from 0.002 to 0.006. For each scenario, 1000 simulations were performed.

ARCHIE calculated cc-values (Figure S19A) that measure the *trans* association between each variant and gene sets across 1000 simulations and scenarios. ARCHIE also selected genes (Figure S19B) from the 101 input genes to be target genes in ARCHIE components. While only 30% of genes have true *trans* effects, ARCHIE selected nearly all 101 genes in the ARCHIE component (Figure S19B). To calculate the p-value of a cc-value, we simulated an empirical null distribution of cc-values by simulating one million null z-scores. Then, an empirical p-value is calculated to be the expected number of null cc-values larger than the observed cc-value (Figure S19C). We found  that ARCHIE p-values are deflated across genetic variances (Figure S19C). To calculate power, p-values were adjusted for multiple testing to control the false positive rate using R package ‘qvalue’[9] (FDR < 0.05, Figure S19D).

ARCHIE was not able to identify significant tests in any of 1000 simulations in our simulation settings, even at the largest genetic variance of 0.006 (Figure 19D). This result is not surprising, as ARCHIE is not designed to detect the type of *trans* signal trans-PCO is designed to detect. Trans-PCO is designed to identify weak *trans* effects from a single SNP to a set of genes, for example, from a *cis*-eQTL of a transcription factor gene to multiple co-regulated genes; therefore, in our simulation settings, one SNP has weak *trans* effects on multiple genes with co-regulated expressions. However, ARCHIE is designed  to detect *trans* effects where multiple SNPs have weak effects on one gene (for example, multiple GWAS variants have weak *trans* effects on a single gene) or more complicated *trans* effects between multiple SNPs and multiple genes (see Figure 2, Dutta et al.[8]).

Dutta et al.[8] performed simulations to evaluate the power of ARCHIE in simulation settings that ARCHIE is designed for. More specifically, in contrast to our simulation settings, ARCHIE simulations simulated multiple SNPs to have weak *trans* effects on a single gene (the simple model in Figure 2 of Dutta et al.[8]) and a more complex model where multiple SNPs having weak *trans* effects on multiple genes (the complex model in Figure 2 of Dutta et al.[8]). However, the simulation was complicated and was not easy to replicate without the original source code (which is not publicly available). To perform fair comparisons between the two methods, we seeked alternative approaches to evaluate whether trans-PCO could replicate ARCHIE signals in real data analyses (i.e. eQTLGen dataset), rather than in simulations. The details can be found in the following two sections.

#### Apply trans-PCO to ARCHIE selected gene sets

Another way we used to compare trans-PCO and ARCHIE was to evaluate whether trans-PCO can identify the *trans* signals identified by ARCHIE in the eQTLGen dataset.  In eQTLGen, Dutta et al.[8] identified gene sets that are significantly associated with disease–associated variants of 29 traits through *trans* regulation, though only results for three traits were publicly available. Specifically, 2 (resp. 1 and 2) selected gene sets were identified to have significant *trans* associations with prostate cancer (resp. schizophrenia and ulcerative colitis)–associated variants, respectively (Figure S19H). Each component contains a set of variants and a set of genes that are associated through *trans* association. Since trans-PCO is designed to identify *trans*-eQTLs of user-specified gene sets, we applied trans-PCO to gene sets in the five components, and identified the genetic variants associated with the gene sets. We then compared the genetic variants identified by trans-PCO to those by ARCHIE. The goal is to check if trans-PCO could replicate ARCHIE signals.

We applied trans-PCO to eQTLGen summary statistics and used ARCHIE-selected gene sets as gene modules. There are five ARCHIE gene sets for three traits (Figure S19). We performed trans-PCO on the five gene sets and calculated association p-values across all eQTLGen variants. The procedure of applying summary-statistics–based trans-PCO is described in the previous section. To estimate the correlation matrix for a gene set, we used the sample correlation among genes using independent null variants, which are defined to have p-values smaller than 1e-4 for all genes in the set (Figure S19E). The ratio between the number of null SNPs and number of genes is high (minimum ratio=53), indicating the estimation of the correlation matrix should be accurate enough to avoid false positive inflation (Figure S8 and Figure S17). We test only *trans* variants that are either more than 5Mb away from genes in the set or on different chromosomes, to be consistent to the definition used in Dutta et al.[8]. P-values were adjusted for multiple testing to control the false positive rate by Bonferroni correction (FDR < 0.05, Figure S19F, Figure S19G).

Among five components, four had significant *trans* associations by trans-PCO with at least one of ARCHIE selected variants in the same component (Figure S19G). Therefore, trans-PCO replicated 80% of the *trans* signals identified by ARCHIE at 5% FDR. In total, ARCHIE identified 134 variants in the five components. Trans-PCO replicated a large proportion (min:36.5%~max:85.1%) of the variants for component 1’s (C1’s) for the three diseases (Figure S19G). Only one out of the 13 variants (7.7%) were replicated by trans-PCO for component 2 (C2) of prostate cancer. Nonetheless, trans-PCO identified 1655 additional significant *trans*-eQTL SNPs at 5% FDR ranging from 65 to 640 for each set (Figure S19F).

In summary, by applying trans-PCO to the five ARCHIE identified gene sets in eQTLGen, we identified 1702 *trans*-eQTL SNPs. ARCHIE identified 134 variants, and 47 (35%) variants are common to both methods. Trans-PCO replicated at least one variant for the corresponding gene set in four out of the five components (Figure S19G). We also found trans-PCO has a better replication of variants in C1’s than C2’s (36.5%-85.1% in C1 vs. 0%-7.7% in C2).

#### Comparison of eQTLGen signals

We compared trans-PCO and ARCHIE by directly comparing the eQTLGen signals detected by the two methods. ARCHIE identified gene sets that are significantly associated with disease–associated variants of 29 traits, though only results for three traits were publicly available. There were five significant ARCHIE components for the three traits. Each component contains a set of SNPs and a set of genes, and the set of SNPs are correlated to the set of genes through *trans* regulation.

We also applied trans-PCO to eQTLGen summary statistics as described in the previous section. We analyzed 129 co-expression gene modules and identified 8116 significant *trans*-eSNP–gene co-expression module pairs, corresponding to 2161 eQTLGen test SNPs and 122 gene modules.

To check if ARCHIE signal components found in eQTLGen were replicated by trans-PCO, we checked (1) if ARCHIE selected genes are included in the significant *trans* target gene modules identified by trans-PCO (Figure S19H), and (2) if ARCHIE selected variants are replicated as *trans*-eQTL SNPs by trans-PCO (Figure S19I). Among selected genes by ARCHIE, all of those included in our eQTLGen analysis were included in a significant *trans*-eQTL module by trans-PCO (Figure S19H). Among selected variants by ARCHIE, 31% (40 out of 129 included variants) were also replicated as significant *trans* signals by trans-PCO (Figure S19I). We note that failing to replicate the remaining ARCHIE signals does not indicate poor performance of trans-PCO for detecting *trans*-eQTLs, as trans-PCO detected 15x more *trans*-eQTL SNPs than ARCHIE. ARCHIE and trans-PCO are designed to detect different *trans* signals: ARCHIE is designed to identify *trans* signals in which a target gene has weak *trans* effects from multiple SNPs (for example from multiple GWAS SNPs to a single gene, as modeled in ARCHIE simulations, Figure 2 of Dutta et al.[8]), whereas trans-PCO is designed to detect *trans* effects from one SNP to multiple genes (for example *trans* effects from a master regulator to multiple downstream genes).

In summary, we compared trans-PCO to ARCHIE in both simulations and in real data analyses. While both are more powerful than the univariate *trans*-eQTL method (Figure 2 of our main text and Figure 2 of Dutta et al.[8]), trans-PCO and ARCHIE are designed to detect different types of weak *trans*-eQTL signals. Our comparison results also support that trans-PCO and ARCHIE are powered at detecting different *trans*-eQTL signals. For example, ARCHIE has no power to detect weak *trans* effects from one SNP to multiple genes in the simulation analyses; while trans-PCO detects a lot more *trans*-eQTL SNPs for the same gene sets than ARCHIE, it only replicate part of the *trans*-eQTL SNPs selected by ARCHIE. There are also other differences between ARCHIE and trans-PCO (see Discussion in the main text), for example, ARCHIE signals are disease specific and the main goal is to identity *trans* genes associated disease associated variants, whereas trans-PCO signals are not disease specific and can be used to perform genome-wide scans of *trans*-eQTLs and produce comprehensive catalogs of *trans*-eQTLs in various tissues and cell types. trans-PCO can be applied to identify *trans*-eQTL SNPs of any user-defined gene sets; in contrast, ARCHIE takes all genes as input and infers a subset of genes *trans* regulated by the variants.

### Compare trans-PCO and Rotival et al.

We compared trans-PCO to the method proposed in Rotival et al.[10], which is to identify *trans*-eQTLs of co-expressed gene sets, and shares the same goal as trans-PCO. Therefore, we used simulations to demonstrate the Rotival et al. method has minimal power to identify weak *trans* effects. We will first describe how the Rotival et al. method works and then demonstrate the performance of the method in simulations.

The Rotival et al. method consists of four main steps: (1) Independent Component Analyses (ICA), a matrix factorisation method, is used to infer components representing co-expression patterns from the expression of all genes; (2) ICA components are then tested against all SNPs to filter out non-suggestive associations (p value>1e-7); (3) for the remaining components and SNPs, a subset of genes contributing strongly to each component are selected as a gene module; (4) for a pair of gene module and a SNP, a significant association is identified if the genes in the module are enriched in genes that are individually associated to the SNP (univariate p-value<1e-5) compared to all other background genes outside the module. The enrichment is tested using the hypergeometric test.

We want to note two points in the comparison of Rotival et al. method and trans-PCO. First, filtering out non-suggestive associations in step (2) can lead to loss of power for identifying *trans*-eQTLs. Second, enrichment analysis to quantify the associations between a gene module and a SNP has limited power at detecting weak *trans* signals. We elaborate our points as follows.

First, we note that filtering associations of ICA components and SNPs in step 2 can be a major power limiting step. To calculate the association between an ICA component and a SNP, the factor loadings of the ICA component are used as the component (or module) profile. As observed in Kolberg et al.[7], the factor loadings of matrix factorisation (or eigengenes) is highly correlated with the first primary PC (PC1) of gene modules defined by co-expression clustering analysis. However, we and others have shown that PC1 has very limited power for detecting *trans* genetic effects between co-regulated gene sets and variants (see more discussions on PC1 having limited power in the comparison with PC1). Therefore, many *trans* signals would have weak associations with PC1, and thus would be removed from the remaining signals used in following steps to identify final *trans* signals.

Additionally, we note that the enrichment analysis by hypergeometric test is less powerful at detecting weak *trans* effects. As stated above, Rotival et al. essentially uses hypergeometric tests to identify *trans*-eQTL SNP-component associations, which are expected to have an enrichment of weak *trans* effects. We therefore performed simulations to evaluate the performance of the enrichment test used by Rotival et al., assuming genes representing co-expression module are already known. The simulations were adapted from the original simulations evaluating the power of trans-PCO as described in Methods ("Simulation"). We first simulated the z-scores between a SNP and $K = 101$ genes in a gene module, following the distribution $N_K(\sqrt{n}\beta, \Sigma_{K\times K})$, where $n$ is sample size (*N* = 500), $\beta$ is a vector representing the true effect sizes of the SNP on $K$ genes and $\Sigma_{K\times K}$is the residualized expression correlation matrix of 101 genes from a real gene module of DGN dataset. Among $K$ genes, a proportion $\gamma$ of them are causal with non-zero effects. Therefore, we generated $\beta_k$ from a point normal distribution, where $\beta_k \sim N(0, \sigma^2{}_b)$ for proportion $\gamma$ ($\gamma = 1\%, 5\%, 10\%, 30\%\ and\ 50\%$), and $\beta_k = 0$, otherwise. The *trans* genetic variance $\sigma^2{}_b$ is set to be 0.001 as default. We also tried larger variances, 0.01, 0.05, 0.1, and 0.2. We simulated 10k SNPs for each simulation and 1k simulations.

To check if target causal genes are enriched in genes included in the module, we also simulated the "background" genes, i.e. genes outside the module. We assume genes outside the module are independent and there are no target causal genes. We simulated 12,001 background genes (12,102 genes used in DGN dataset, subtracted by 101 genes included in the module) from standard normal distribution with zero effects. To define significant individual associations for enrichment, we used p-value cutoff 1e-5 to be consistent with Rotival et al.. Then the enrichment p-value of a SNP for the gene module was calculated by the hypergeometric test. P-values were adjusted using 'qvalue' at FDR<0.1. Power was calculated as the proportion of SNPs that were identified to be significant among 10k SNPs.

As shown in Figure S20, Rotival et al. has minimal power at detecting trans associations in the case of weak effects. Under the setting where genetic variance is 0.001, the enrichment test has no power across all causal proportions, while trans-PCO has much higher power, for example, power is 37% when 30% genes are true target genes. We note that the enrichment test can have power for detecting *trans*-eQTLs when the *trans* effects are large (which is not common for trans-eQTLs). For example, when genetic variance is 0.01, the enrichment test has a power of 32% when 5% genes are target causal genes. Trans-PCO has a higher power of 64% under the same setting. In the case of even larger genetic variances, e.g. 0.1 and 0.2, the enrichment test has a comparable power with trans-PCO. In summary, the enrichment test does not have power to detect multiple weak *trans* effects.

#### Drug targets are associated with immune-related gene sets in *trans*

To show the translational application of trans-PCO results, we examined whether drug targets are more likely to be associated with disease-relevant pathways or gene sets in *trans*. We first downloaded drug targets of various diseases from The Broad Institute Drug Repurposing Hub (https://repo-hub.broadinstitute.org/repurposing). We focused on the disease allergy, because it is immune-related given our analyzed gene expression datasets are from blood tissue. It has a relatively large number of drug target genes (55 launched targets), 5 of which are near (within 1Mb) allergy associated SNPs in eQTLGen (~10k SNPs used for *trans* analysis). We identified SNPs that are significantly associated with allergy using allergy GWAS summary statistics[11] (Table S8, p-value<5e-8). We then examined whether these 5 drug targets are near any SNPs that have significant *trans* associations with immune-related gene co-expression modules or hallmark gene sets in the eQTLGen dataset. Among a total of 142 gene sets (129 co-expression gene modules and 11 hallmark gene sets) used in eQTLGen analysis, 19 were defined as immune-related. Interestingly, we found that all 5 drug target genes near allergy loci are associated with an immune-related gene set through *trans* regulation. Details of the targets and their associated immune-relevant gene sets can be found in Table S19. While the enrichment of allergy drug targets in *trans*-eQTLs of immune-related gene sets is not statistically significant (Table S20), it is likely due to the small number of drug targets in the analyses. Additionally, it is encouraging to see that the gene sets associated with the drug targets are highly relevant to allergy, for example, B cell receptor signaling pathway is associated with three of the drug targets.

### Supplementary Figures

Figure S1. Simulation results at various genetic variances, including at extremely low proportions of causal genes.

Figure S2. Quantile-quantile plot of P values from null simulations.

Figure S3. Colocalization of *trans*-eQTLs and *cis*-eQTLs at (A) *NFE2* and (B) *PLAGL1* loci.

Figure S4. Gene ontology enrichment of co-expression gene modules (A) M3 and (B) M4.

Figure S5. 965 *trans*-eSNP-module pairs in DGN associated with 50 MSigDB hallmark gene sets representing well-defined biological processes.

Figure S6. Colocalization of *trans*-eQTLs of the heme metabolism and various red blood traits.

Figure S7. Heritability enrichment of gene module M3 in blood traits estimated by S-LDSC.

Figure S8. Summary-statistic–based trans-PCO is well controlled for test statistics inflation.

Figure S9. The trans-PCO P values in eQTLGen are much smaller than in DGN.

Figure S10. Associations at the *ARHGEF3* locus with gene modules in both DGN and eQTLGen.

Figure S11. Distribution of eigenvalues of gene module 1.

Figure S12. P value cutoff to define null SNPs for gene modules.

Figure S13. Comparison between *trans*-eQTLs detected by trans-PCO and PC1 methods.

Figure S14. Simulation scenario when PC1 has the highest power.

Figure S15. Trans-PCO analyses of co-expression gene modules in DGN.

Figure S16. Heritability enrichment of all gene modules in all traits.

Figure S17. Ratio of independent null SNPs over module size across 50 MSigDB biological processes.

Figure S18. Simulation scenario when parameters are large.

Figure S19. Comparison of trans-PCO and ARCHIE.

Figure S20. Comparison of trans-PCO and Rotival et al..

464

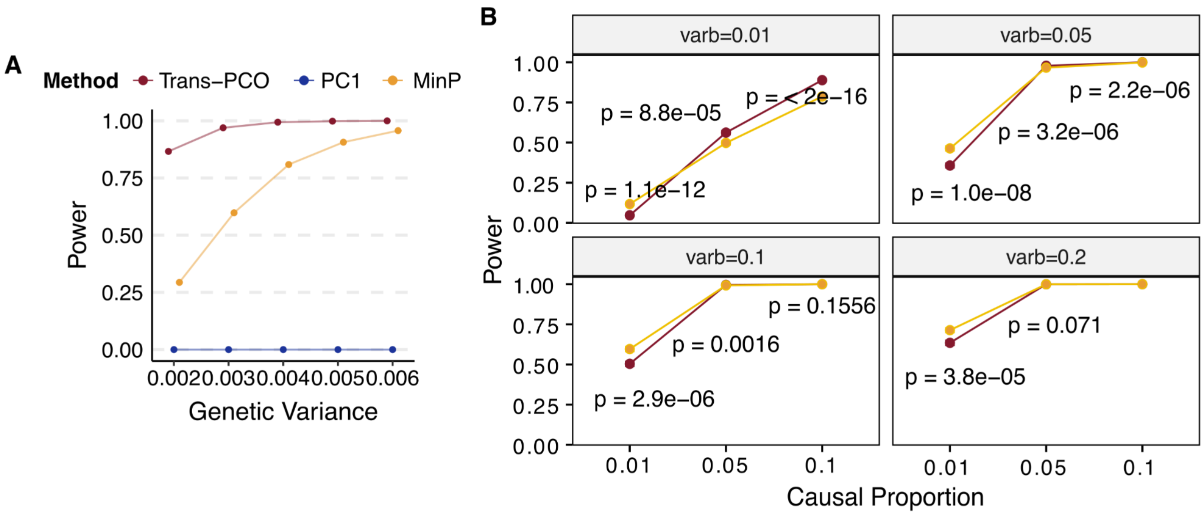


**Figure S1. Simulation results at various genetic variances. (A)** We compared the power of trans-PCO, PC1, and MinP methods under genetic variances 0.002, 0.003, 0.004, 0.005, 0.006. We used the same gene module as in Figure 2. We simulated sample size to be 500 and the proportion of target genes with non-zero effects in the gene module to be 30%. Power was computed from 10k SNPs across 1000 simulations. The error bars are 95% confidence intervals. Many are too small to be visible. **(B) Various genetic variances at extremely low proportions of causal genes. Simulation scenarios when univariate test can be more powerful than multivariate test trans-PCO.** We compared the power of trans-PCO and MinP methods under large effect sizes with high levels of sparsity. Specifically, we simulated the proportion of target genes with non-zero effects to be 1%, 5%, and 10%, and large genetic variances to be 0.01, 0.05, 0.1, and 0.2. We used the same gene module as in Figure 2 and simulated the sample size to be 500. Power was computed from 10k SNPs across 1000 simulations. P-values are from the Wilcoxon test to compare two group means. We observe that univariate method (“MinP”) has significantly higher power than multivariate method (“trans-PCO”) when the sparsity level is high and effect sizes are large. For example, in the case of large genetic variance (varb=0.05) and one gene being causal (casual proportion 1%), MinP has significantly higher power than trans-PCO (p-value=1e-8). As causal genes increase, both MinP and trans-PCO have increased power. To be noted, in the case of small genetic variance (varb=0.01) with weak effects, trans-PCO gains more power than MinP as it aggregates multiple weak effects to improve power.

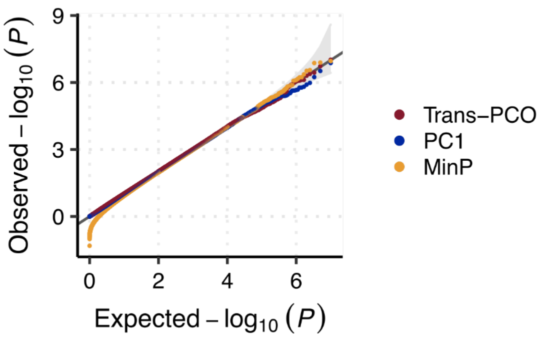


**Figure S2. Quantile-quantile plot of P values from null simulations.** We simulated 10 million null SNPs with zero effects for genes in the simulated gene module (same as Figure 2 and Figure S1). We tested the *trans* association of simulated SNPs with gene modules using trans-PCO, PC1, and MinP methods and calculated the P values. Colors represent different methods.

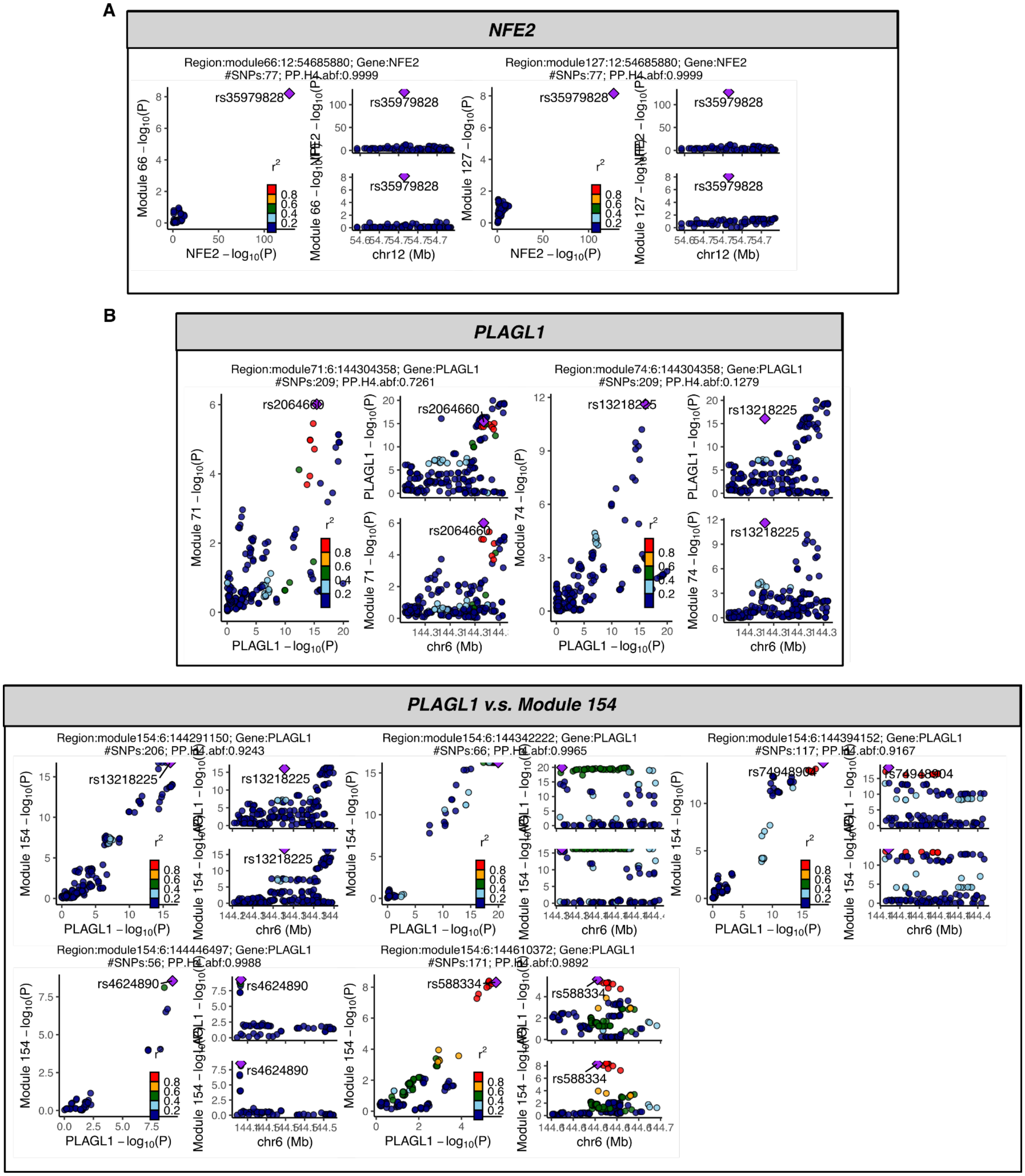


**Figure S3. Colocalization of *trans*-eQTLs and *cis*-eQTLs at (A) *NFE2* and (B) *PLAGL1* loci.** Each sub plot represents a genomic region that has a shared causal variant for the corresponding *trans* target gene module and *cis* gene. The plot title gives (1) the coloc region, which is defined as the *trans* target gene module and the lead *trans*-eQTL in this region, (2) the *cis* gene near the *trans*-eQTL, (3) the number of SNPs in the region, (4) PP4.

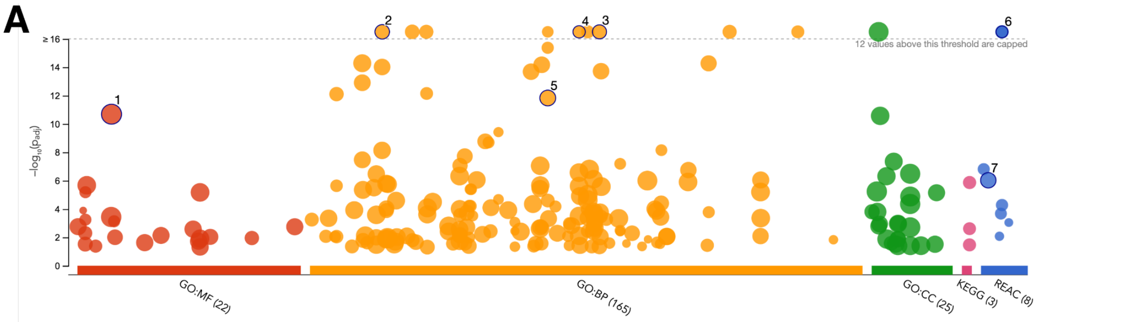


| ID | Source | Term ID | Term Name | $p_{adj}$ (query_1) |
|---|---|---|---|---|
| 1 | GO:MF | GO:0005515 | protein binding | $2.039\times10^{-11}$ |
| 2 | GO:BP | GO:0009615 | response to virus | $3.176\times10^{-27}$ |
| 3 | GO:BP | GO:0051607 | defense response to virus | $1.413\times10^{-30}$ |
| 4 | GO:BP | GO:0048525 | negative regulation of viral process | $9.596\times10^{-18}$ |
| 5 | GO:BP | GO:0045087 | innate immune response | $1.469\times10^{-12}$ |
| 6 | REAC | REAC:R-HSA-9... | Interferon alpha/beta signaling | $4.571\times10^{-18}$ |
| 7 | REAC | REAC:R-HSA-1... | Cytokine Signaling in Immune system | $9.350\times10^{-7}$ |

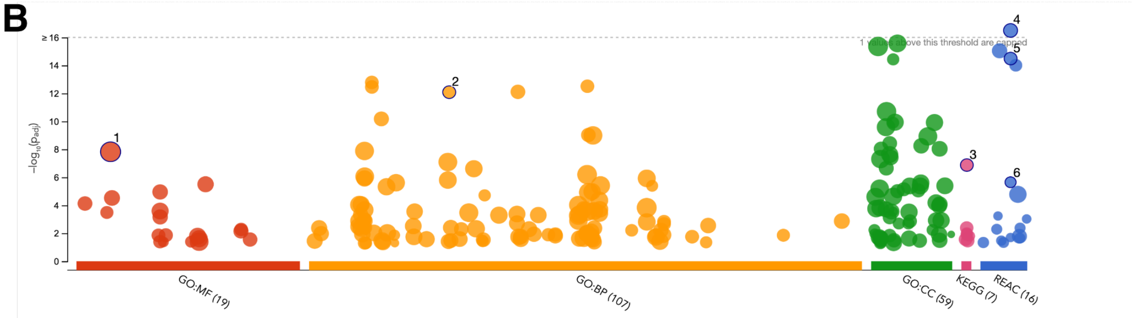


| ID | Source | Term ID | Term Name | $p_{adj}$ (query_1) |
|---|---|---|---|---|
| 1 | GO:MF | GO:0005515 | protein binding | $1.495\times10^{-8}$ |
| 2 | GO:BP | GO:0030168 | platelet activation | $8.163\times10^{-13}$ |
| 3 | KEGG | KEGG:04611 | Platelet activation | $1.328\times10^{-7}$ |
| 4 | REAC | REAC:R-HSA-7... | Platelet activation, signaling and aggregation | $1.062\times10^{-21}$ |
| 5 | REAC | REAC:R-HSA-1... | Platelet degranulation | $3.229\times10^{-15}$ |
| 6 | REAC | REAC:R-HSA-7... | Platelet Aggregation (Plug Formation) | $2.241\times10^{-6}$ |

**Figure S4. Gene ontology enrichment of co-expression gene modules (A) M3 and (B) M4.** We used four term categories to look at the enrichment in gene modules[6], including GO:MF, GO:BP, KEGG, and REAC. Categories are shown in colors. The y-axis shows the adjusted enrichment P values. We highlighted a few most significant and interesting enrichment terms.

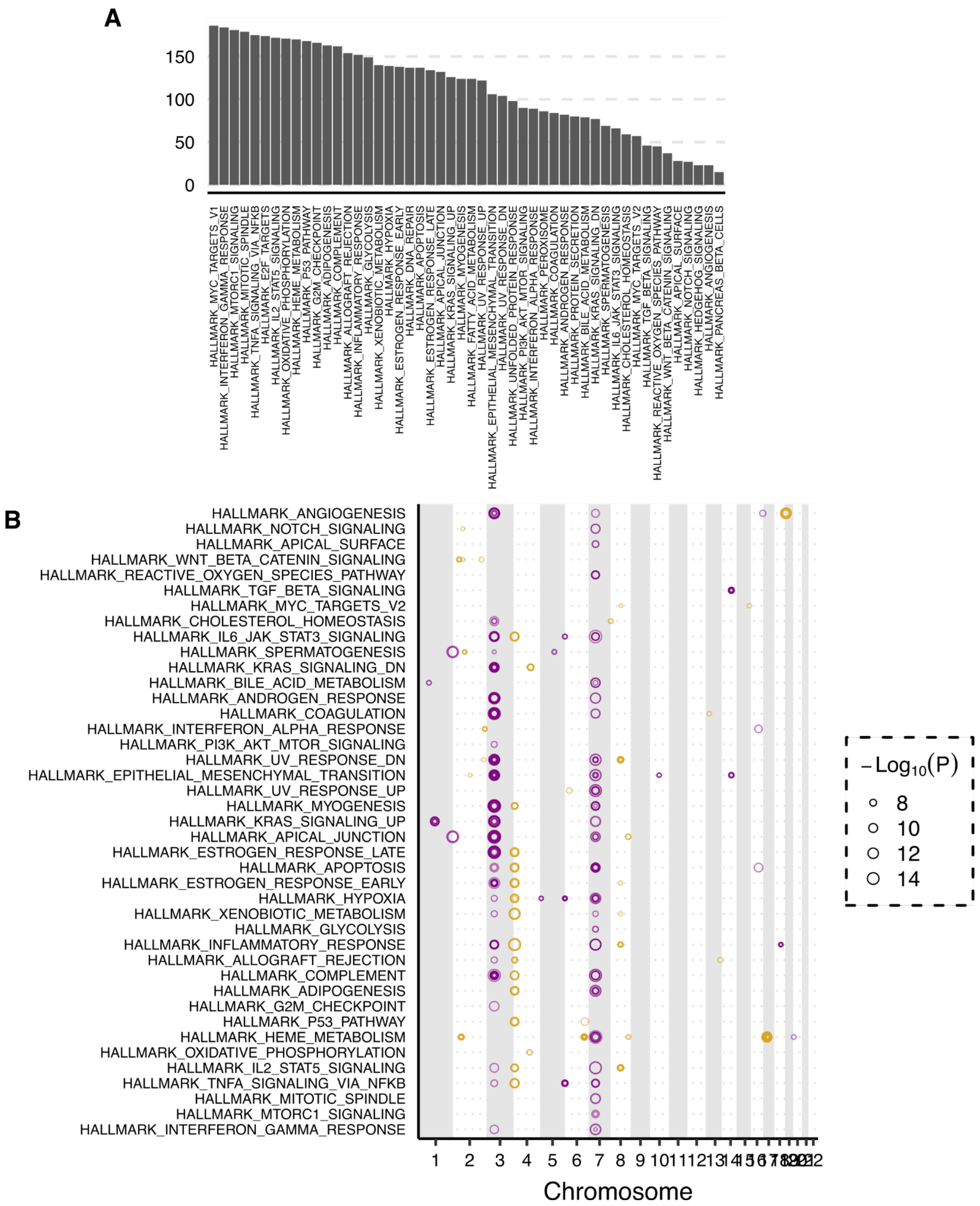


**Figure S5. 965 *trans*-eSNP-module in DGN associated with 50 MSigDB hallmark gene sets representing well-defined biological processes. (A) Number of genes in the hallmark gene sets.** Names of the biological processes are shown on the x-axis. Sizes are shown on the y-axis. **(B) *Trans*-eQTLs of the biological processes.** The gene sets are shown on the y-axis. The chromosome position of their corresponding *trans*-eQTLs is on the x-axis. Colors represent odd and even chromosomes to better distinguish the positions. Point size represents the P value of each pair of (*trans*-eSNP, gene set).

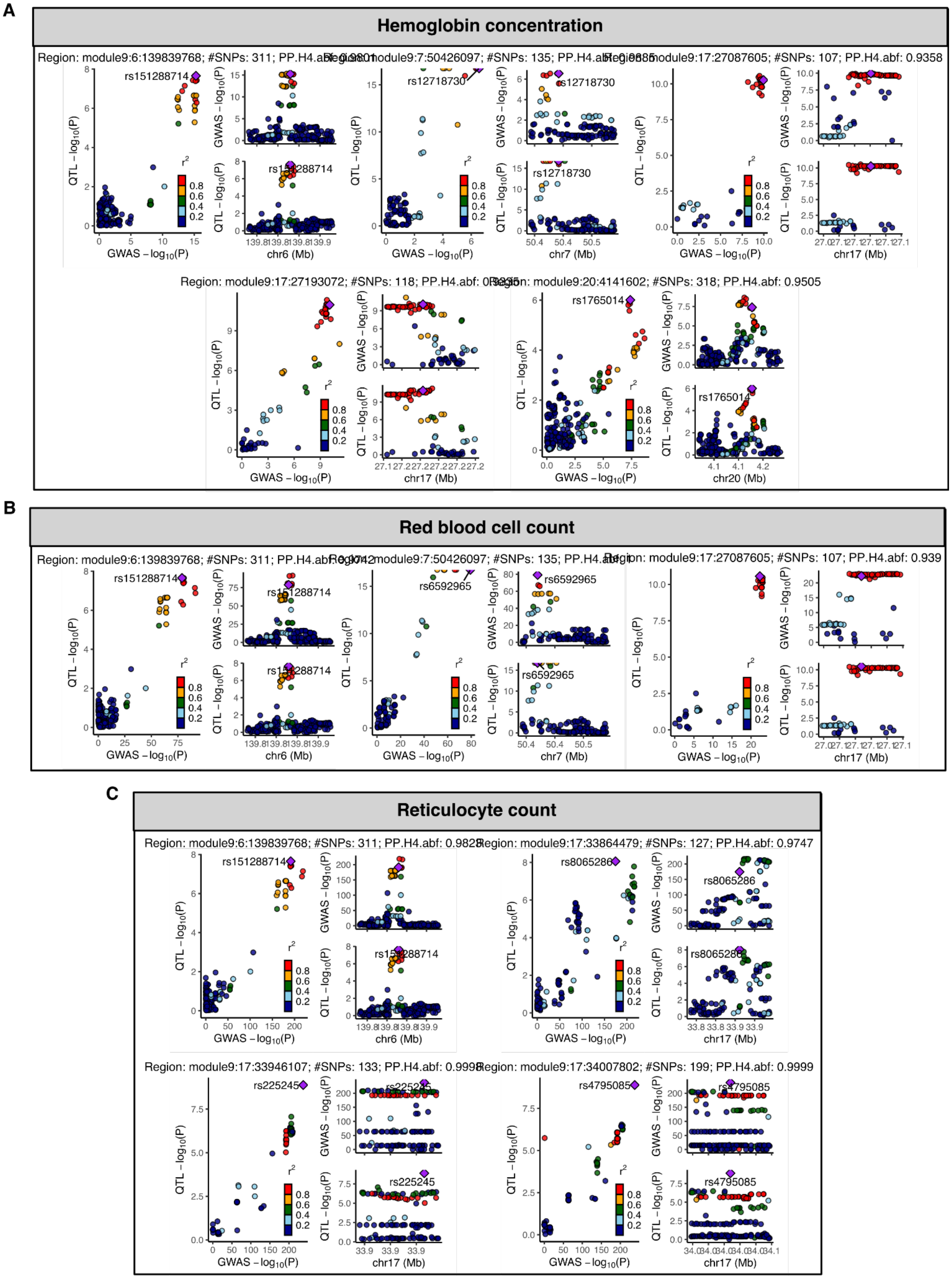
A
Hemoglobin concentration
Region: module9:6:139839768; #SNPs: 311; PP.H4.abf: 0.9801
Region: module9:7:50426097; #SNPs: 135; PP.H4.abf: 0.9885
Region: module9:17:27087605; #SNPs: 107; PP.H4.abf: 0.9358
Region: module9:17:27193072; #SNPs: 118; PP.H4.abf: 0.9355
Region: module9:20:4141602; #SNPs: 318; PP.H4.abf: 0.9505
rs151288714
rs12718730
rs1765014
B
Red blood cell count
Region: module9:6:139839768; #SNPs: 311; PP.H4.abf: 0.9742
Region: module9:7:50426097; #SNPs: 135; PP.H4.abf
Region: module9:17:27087605; #SNPs: 107; PP.H4.abf: 0.939
rs151288714
rs6592965
C
Reticulocyte count
Region: module9:6:139839768; #SNPs: 311; PP.H4.abf: 0.9828
Region: module9:17:33864479; #SNPs: 127; PP.H4.abf: 0.9747
Region: module9:17:33946107; #SNPs: 133; PP.H4.abf: 0.9998
Region: module9:17:34007802; #SNPs: 199; PP.H4.abf: 0.9999
rs151288714
rs8065286
rs225245
rs4795085
QTL − log10(P)
GWAS − log10(P)
chr6 (Mb)
chr7 (Mb)
chr17 (Mb)
chr20 (Mb)
r2
0.8
0.6
0.4
0.2

**Figure S6. Colocalization of *trans*-eQTLs of the heme metabolism and various red blood traits: (A) hemoglobin concentration, (B) red blood cell count, (C) reticulocyte count.** The sub plots show colocalization of *trans*-eQTLs of heme metabolism and GWAS loci. The plot title gives (1) the coloc region, which is defined as the *trans* target gene module and the lead *trans*-eQTL in this region, (2) the number of SNPs in the region, (3) PP4.

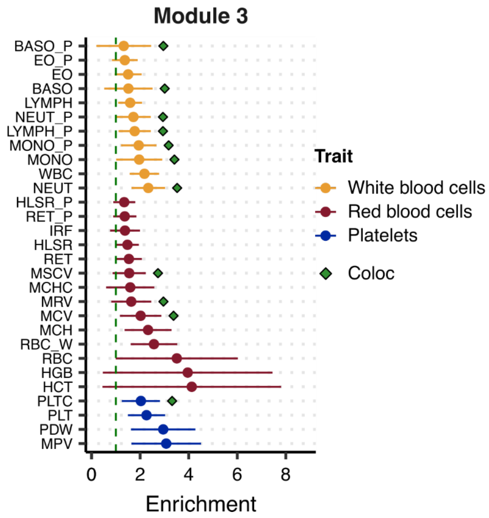


**Figure S7 Heritability enrichment of gene module M3 in blood traits estimated by S-LDSC.** The y-axis shows the blood traits. Colors represent trait types. The heritability enrichment in module 3 is shown on the x-axis. Error bars represent 95% confidence intervals. Green points indicate that there is significant colocalization of the gene module 3 and the trait.

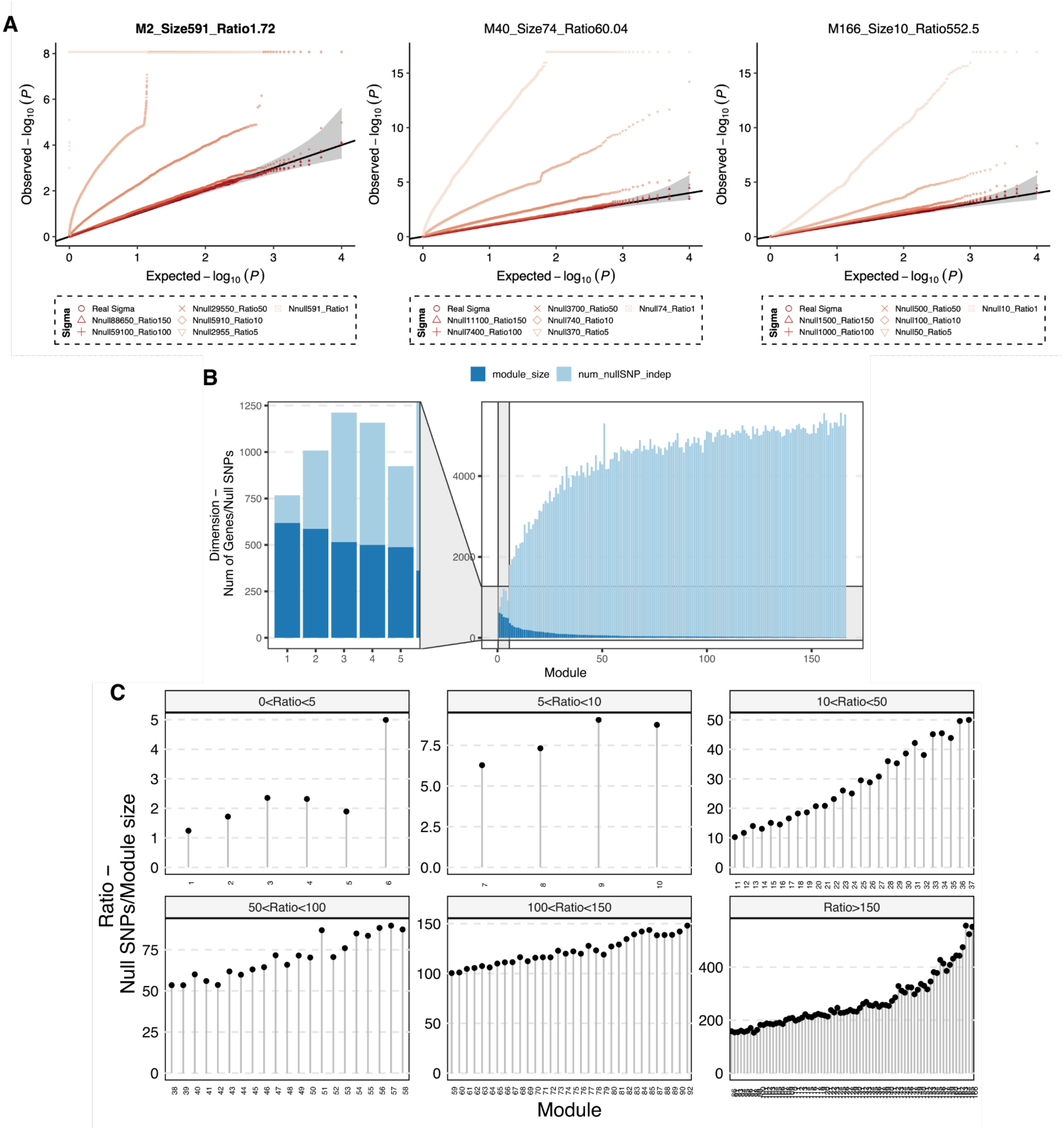
A
M2_Size591_Ratio1.72
M40_Size74_Ratio60.04
M166_Size10_Ratio552.5
Observed – log10 (P)
Expected – log10 (P)
Sigma
Real Sigma
Nnull88650_Ratio150
Nnull59100_Ratio100
Nnull29550_Ratio50
Nnull5910_Ratio10
Nnull2955_Ratio5
Nnull591_Ratio1
Nnull11100_Ratio150
Nnull7400_Ratio100
Nnull3700_Ratio50
Nnull740_Ratio10
Nnull370_Ratio5
Nnull74_Ratio1
Nnull1500_Ratio150
Nnull1000_Ratio100
Nnull500_Ratio50
Nnull100_Ratio10
Nnull50_Ratio5
Nnull10_Ratio1
B
module_size
num_nullSNP_indep
Dimension – Num of Genes/Null SNPs
Module
C
0<Ratio<5
5<Ratio<10
10<Ratio<50
50<Ratio<100
100<Ratio<150
Ratio>150
Ratio – Null SNPs/Module size
Module

**Figure S8. Summary-statistic–based trans-PCO is well controlled for test statistics inflation. (A) Noisy correlation matrices lead to trans-PCO test statistics inflation.** In addition to the gene module 1 in Figure 5A, we also looked at other gene modules with different sizes to investigate how the noisy correlation matrix estimation affects the signal inflation. Here, we show a few more gene modules, including module 2 with size 591, module 40 with size 74, and module 166 with size 10. The correlation estimation by different ratios of null SNPs over the module size is represented by different point shapes and shades. We observe inflated null P values when correlation matrices were less accurately estimated by lower ratios (of null SNPs over module size). **(B) Size of co-expression gene modules and the number of null SNPs used to estimate the correlation matrix.** Light blue bar shows the module size. Dark blue bar shows the number of independent null SNPs found in eQTLGen that were used to estimate the correlation matrix of the gene module. **(C) Ratio of null SNPs over module size across all co-expression gene modules.**

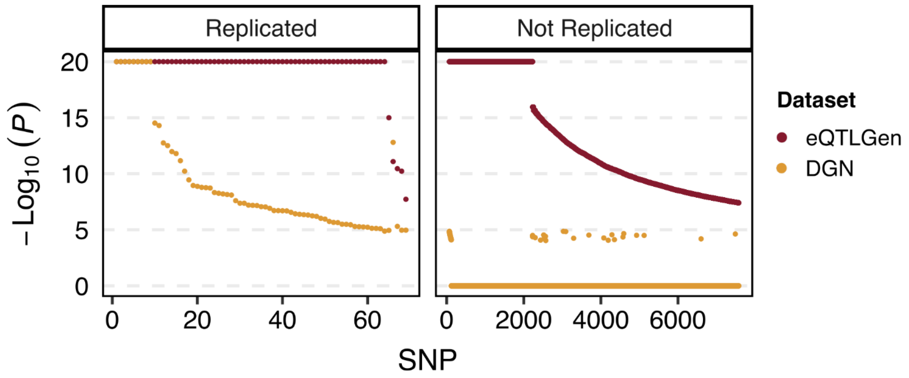


**Figure S9. The trans-PCO P values in eQTLGen are much smaller than in DGN.** We compared the P value of the same pair of SNP and gene module in DGN and eQTLGen. SNP-module pairs are shown on the x-axis. P values are on the y-axis. DGN and eQTLGen P values are colored in yellow and red, respectively. We divided the SNPs into two categories, (1) SNPs identified as *trans*-eQTLs in both DGN and eQTLGen (left panel, “Replicated”), (2) SNPs identified as *trans*-eQTLs only in eQTLGen not in DGN (right panel, “Not Replicated”). We can observe that, first, trans-PCO P values in eQTLGen are smaller than P values in DGN. Second, the replicated *trans*-QTLs have much smaller P values in DGN than those not replicated.

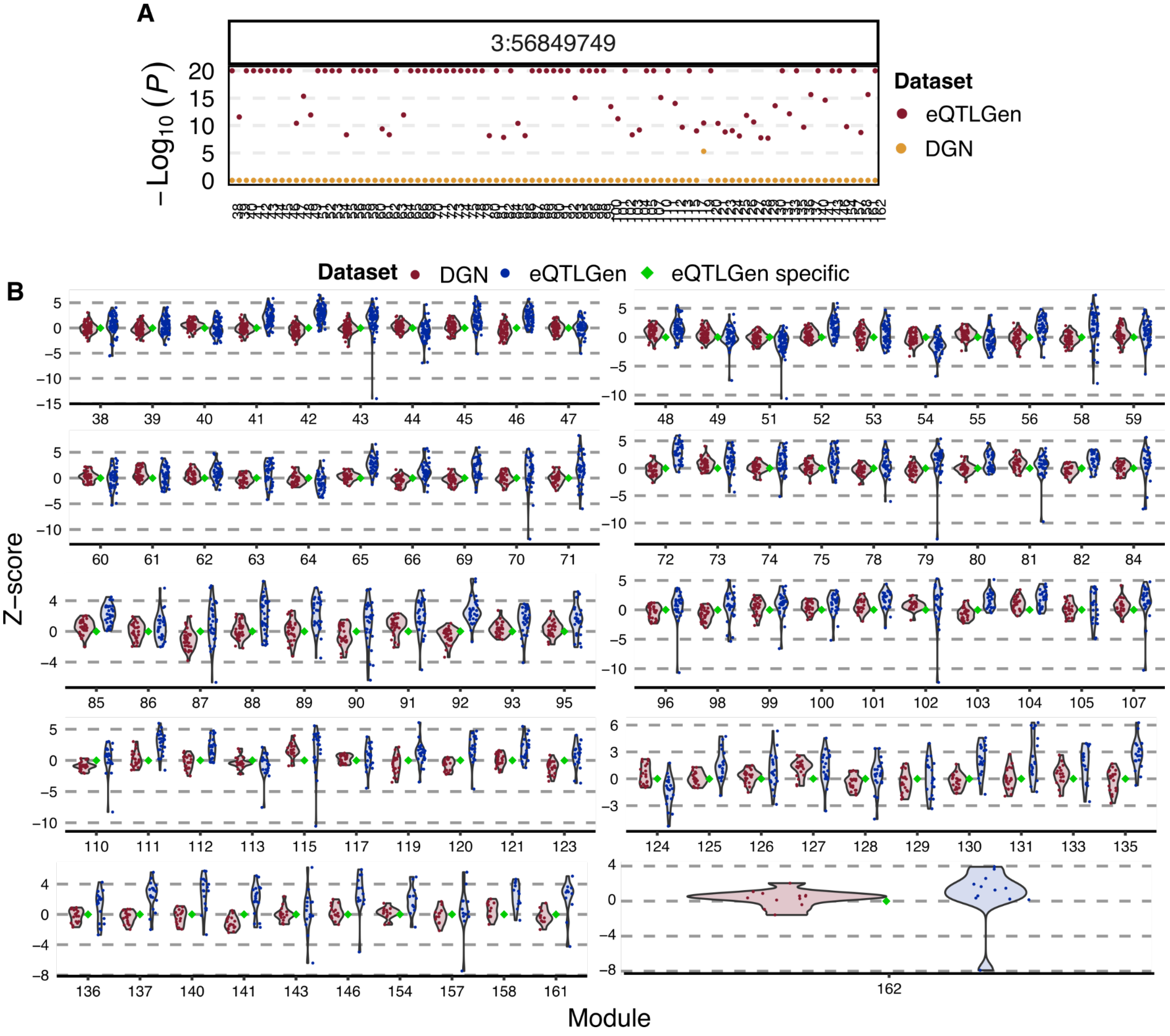


**Figure S10. Associations at the *ARHGEF3* locus with gene modules in both DGN and eQTLGen. (A) Associations P values of SNP 3:56849749 at the *ARHGEF3* locus with gene modules analyzed in eQTLGen.** The Y-axis shows the association P values in eQTLGen (red) and DGN (yellow). Gene modules are on the x-axis. This SNP has much smaller P values in eQTLGen than in DGN. **(B) Z-scores in both DGN and eQTLGen of SNP 3:56849749 across genes in gene modules.** The X-axis shows gene modules. The Y-axis shows z-scores of SNP 3:56849749 with the genes in the corresponding module. DGN and eQTLGen z-scores are shown in red and blue, respectively. Green dot indicates that the SNP and the gene module is a signal specific to eQTLGen.

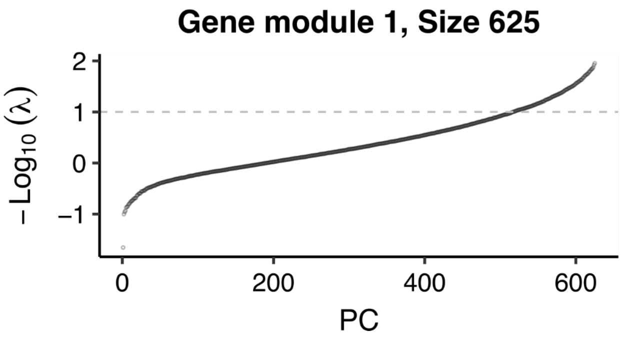


**Figure S11. Distribution of eigenvalues of gene module 1.** We use gene modules 1 as an example. PC's are shown on the x-axis. The eigenvalues are shown on the y-axis. The dashed line represents the eigenvalue cutoff (0.01) we used to define PC's included in the test. We can see the last few PC's have extremely small eigenvalues.

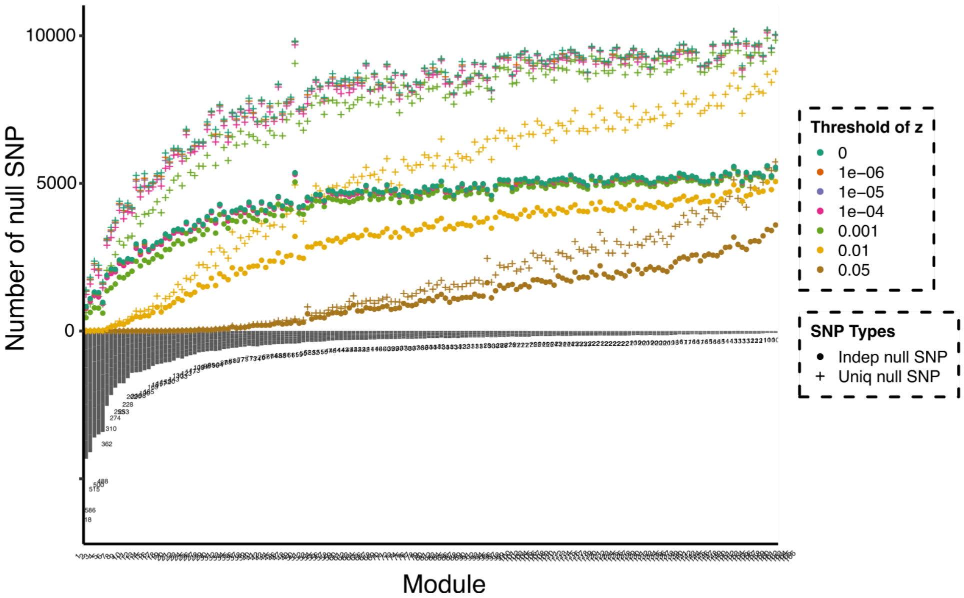


**Figure S12. P value cutoff to define null SNPs for gene modules.** We show the gene modules on the x-axis. The module sizes are shown on the lower y-axis. The upper y-axis represents the number of null SNPs in eQTLGen that are insignificantly associated with all genes in a gene module. Cross shapes represent the null SNPs. Circles represent the LD independent null SNPs ($R2<0.2$). Colors show different p-value cutoffs, 0, 1e-6, 1e-5, 1e-4, 1e-3, 1e-2, and 0.05, to define null SNPs.

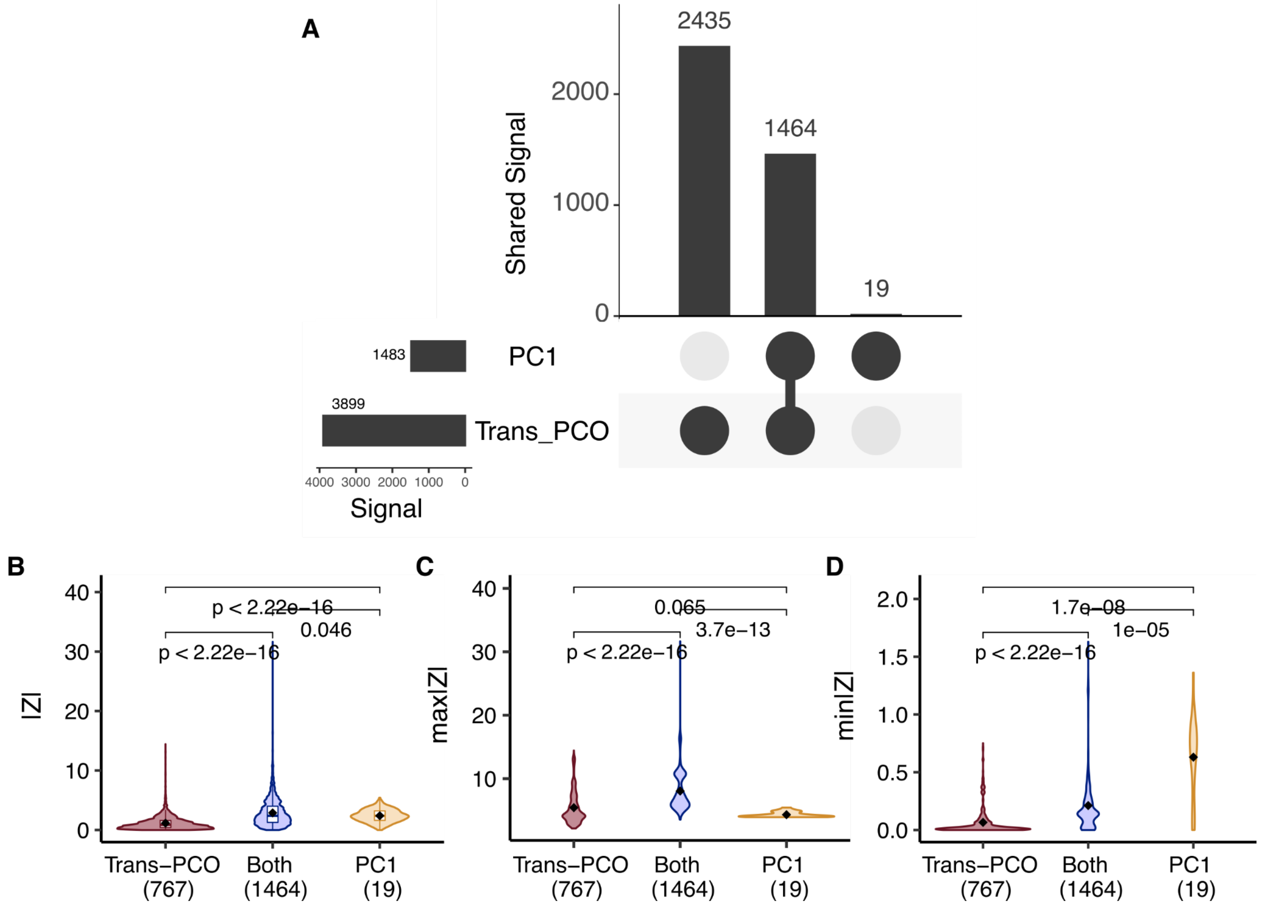


**Figure S13. Comparison between *trans*-eQTLs detected by trans-PCO and PC1[7] methods. (A) Signal comparison of PC1 and trans-PCO.** The first column shows there are 2435 *trans*-eSNP-module signal pairs that are detected only by trans-PCO not by PC1. The second column shows there are 1464 *trans* pairs identified by both trans-PCO and PC1. The last column shows 19 *trans* signals are identified only by PC1. The horizontal bars on the left show the total number of *trans* signals identified by trans-PCO and PC1, respectively. **(B)-(D) Z scores comparison of PC1 and trans-PCO signals.** We divided the *trans*-QTLs into three categories, trans-PCO specific signals in red, trans-PCO and PC1 shared signals in blue, and PC1 specific signals in yellow. We compared the z-scores of the three types of *trans*-eQTLs, in terms of (B) the absolute z-scores of signals for all genes in gene modules, (C) the maximum absolute z-scores of signals across genes in gene modules, (D) the minimum absolute z-scores of signals across genes in gene modules. We can see PC1 specific signals have higher z-scores than signals detected by only trans-PCO or both methods, supporting that trans-PCO can detect much weaker *trans* genetic effects.

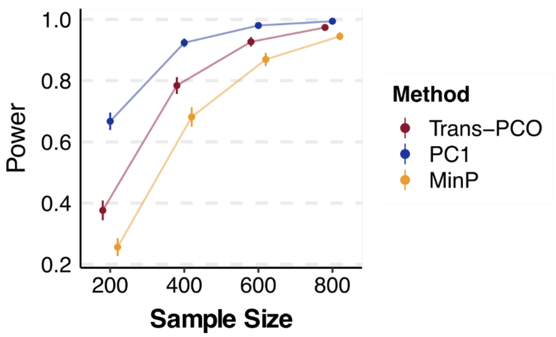


**Figure S14. Simulation scenario when PC1 has the highest power.** As proved in Liu et al.[4], PC1 method performs best in the case where the effects vector of SNP on genes in the module align with the first eigenvector of the gene module. We used the same gene module as in Figure 2. We simulated the effect of SNPs on genes in the module to be $\sqrt{\sigma^2{}_b}\mu_1$, where $\sigma^2{}_b$ is the genetic variance with value 0.001, $\mu_1$ is the first eigenvector of the gene module. We simulated the sample size to be 500, and the proportion of target genes with non-zero effects in the gene module to be 30%. Power was computed from 10k SNPs across 1000 simulations. The error bars are 95% confidence intervals.

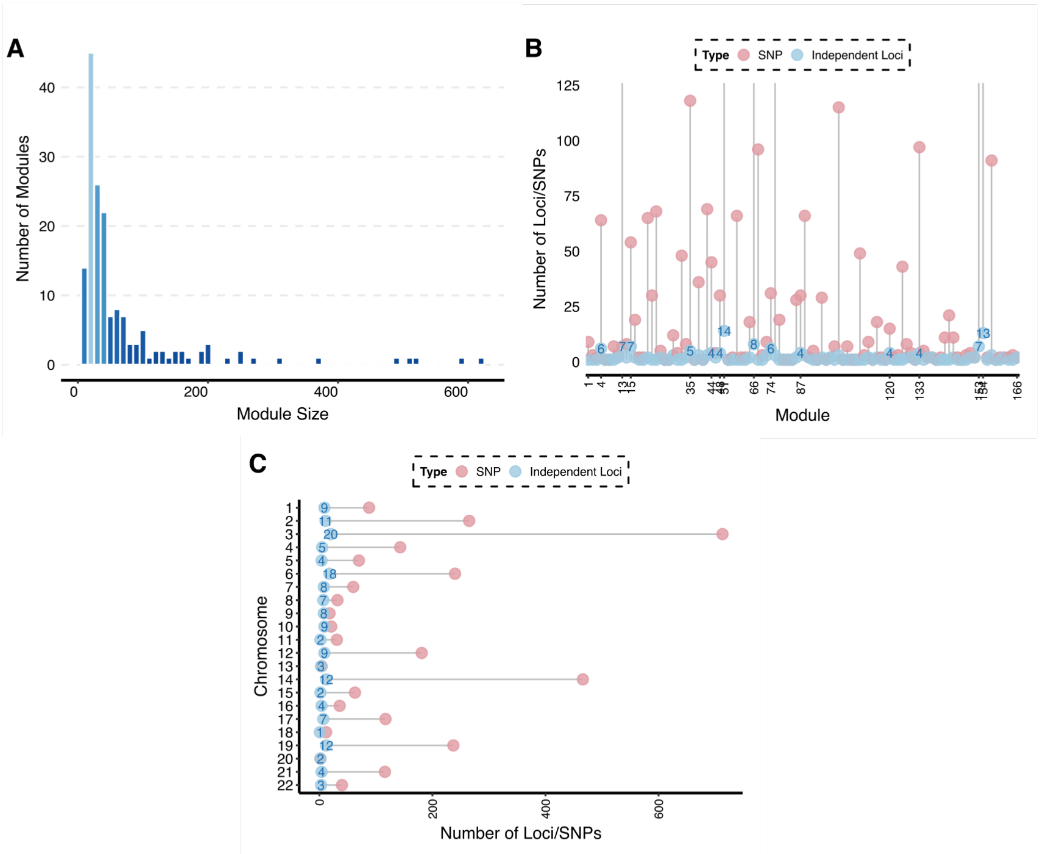


**Figure S15. Trans-PCO analyses of co-expression gene modules in DGN. (A) Size distribution of co-expression gene modules. (B)-(C) The number of *trans*-eQTL signals associated with co-expression modules (B) per module and (C) per chromosome.** Red points represent the number of *trans*-eSNPs. Blue points represent the number of LD independent loci (R2<0.2).

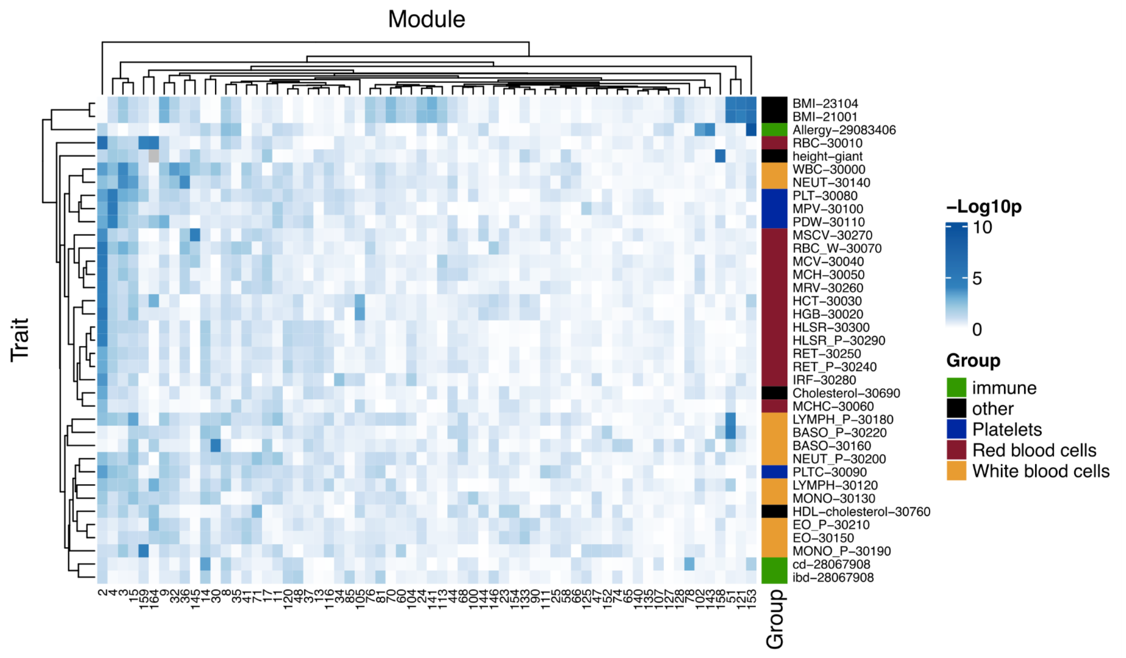


**Figure S16. Heritability enrichment of all gene modules in all traits.** Each row represents a trait. Traits are colored based on the trait types, including white blood cell traits, red blood cell traits, platelet traits, immune diseases, and other traits. Gene modules are shown on the x-axis. We only show the modules that have at least one colocalized region with a trait. Each tile shows the P values of trait heritability enrichments. The rows and columns are clustered based on the heritability enrichment. We can see traits of the same type share similar heritability enrichment patterns.

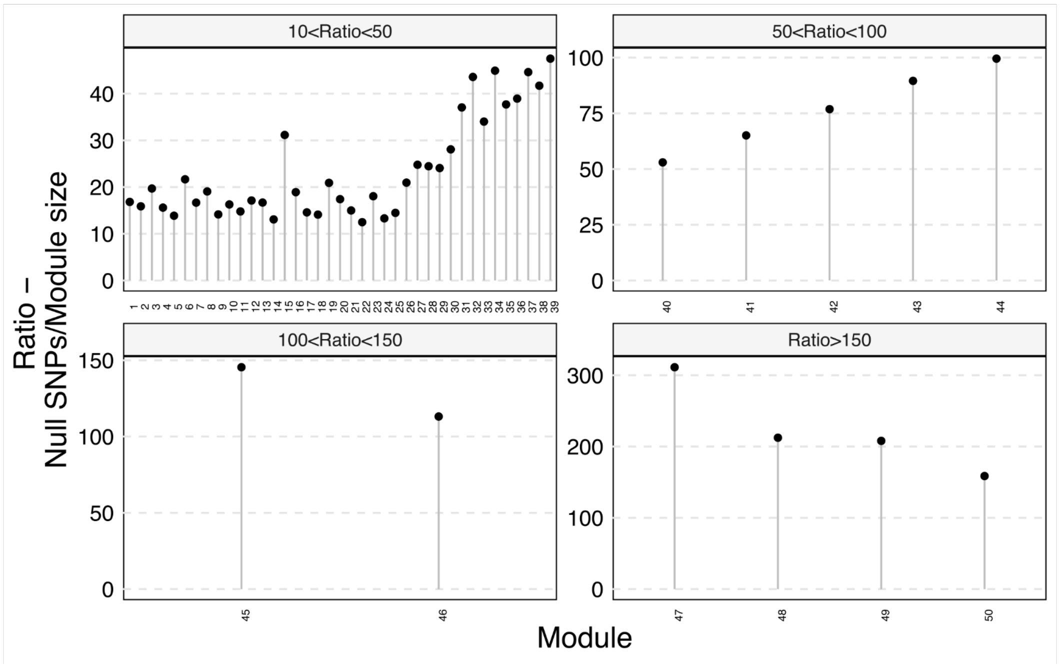


**Figure S17. Ratio of independent null SNPs over module size across 50 MSigDB biological processes.** The gene sets are shown on the x-axis. The ratio is shown on the y-axis. The gene sets are divided into four categories representing different ratios.

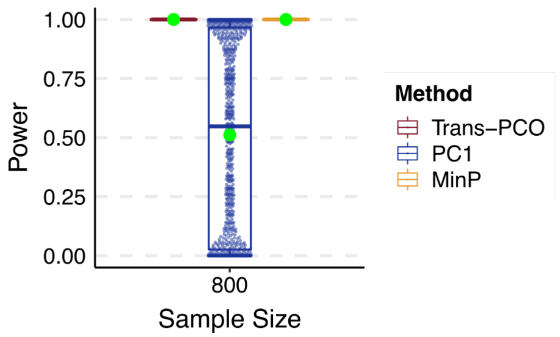


**Figure S18. Simulation scenario when parameters are large.** We used the same gene module as in Figure 2. We simulated the genetic variance to be as large as 0.2, the proportion of target genes with non-zero effects in the gene module to be 100%, and the sample size to 800. Power was computed from 10k SNPs across 1000 simulations. Green points represent the mean power. Each dot represents a simulation.

639

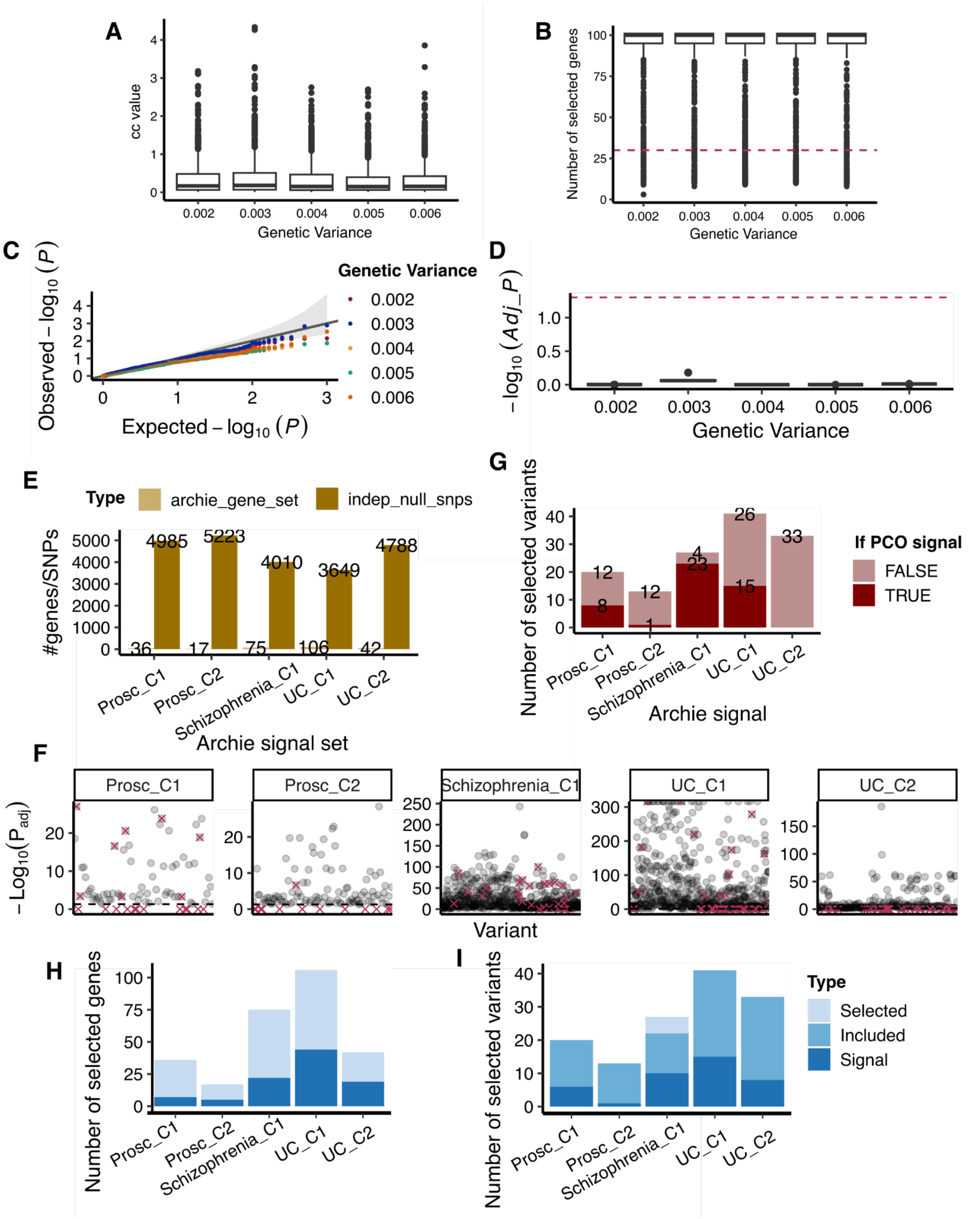

A
cc value
Genetic Variance
0.002
0.003
0.004
0.005
0.006
B
Number of selected genes
C
Observed − $\log_{10}(P)$
Expected − $\log_{10}(P)$
Genetic Variance
D
$-\log_{10}(Adj\_P)$
E
Type
archie_gene_set
indep_null_snps
#genes/SNPs
Archie signal set
G
Number of selected variants
If PCO signal
FALSE
TRUE
Archie signal
F
$-\mathrm{Log}_{10}(P_{adj})$
Prosc_C1
Prosc_C2
Schizophrenia_C1
UC_C1
UC_C2
Variant
H
Number of selected genes
I
Number of selected variants
Type
Selected
Included
Signal

**Figure S19. Comparison of trans-PCO and ARCHIE in Dutta et al.[8]. (A)-(D) Simulation comparison.** (A) Distribution of ARCHIE cc-values across various genetic variances. (B) Number of selected genes by ARCHIE across various genetic variances. Red line indicates the number of true target genes, i.e. 30. (C) QQ plot of empirical p-values across various genetic variances. (D) Distribution of adjusted p-values across various genetic variances. Red line shows FDR level 0.05. **(E)-(G) Trans-PCO on ARCHIE selected gene sets.** (E) X-axis shows significant ARCHIE components for three traits, prostate cancer (Prosc), Schizophrenia, and Ulcerative Colitis (UC). C1 and C2 mean the first and second component. Each component is a pair of selected genes set and variants set. Y-axis shows the size of ARCHIE gene sets (light) and the number of independent null variants (dark) used to estimate correlation matrix of the gene sets. (F) P-values (with Bonferroni correction) of each variant and ARCHIE gene set by trans-PCO. Each panel is an ARCHIE component. Red cross indicates variants selected by ARCHIE. Grey line shows FDR level 0.05. (G) Number of ARCHIE variants across components that are also significant by trans-PCO (dark). **(H)-(I) Comparison of eQTLGen signals by trans-PCO and ARCHIE.** (H) Number of selected genes across ARCHIE components. Lightest blue (Selected) represents the number of selected genes by ARCHIE of each component. Darker blue (Included) represents the selected genes included in trans-PCO eQTLGen analysis. Darkest (Signal) represents genes included in a significant *trans* target gene module by trans-PCO. All ARCHIE selected genes that are analyzed by trans-PCO are in significant *trans* gene modules by trans-PCO. (I) Number of selected variants across ARCHIE components. Labels are similar as in (H).

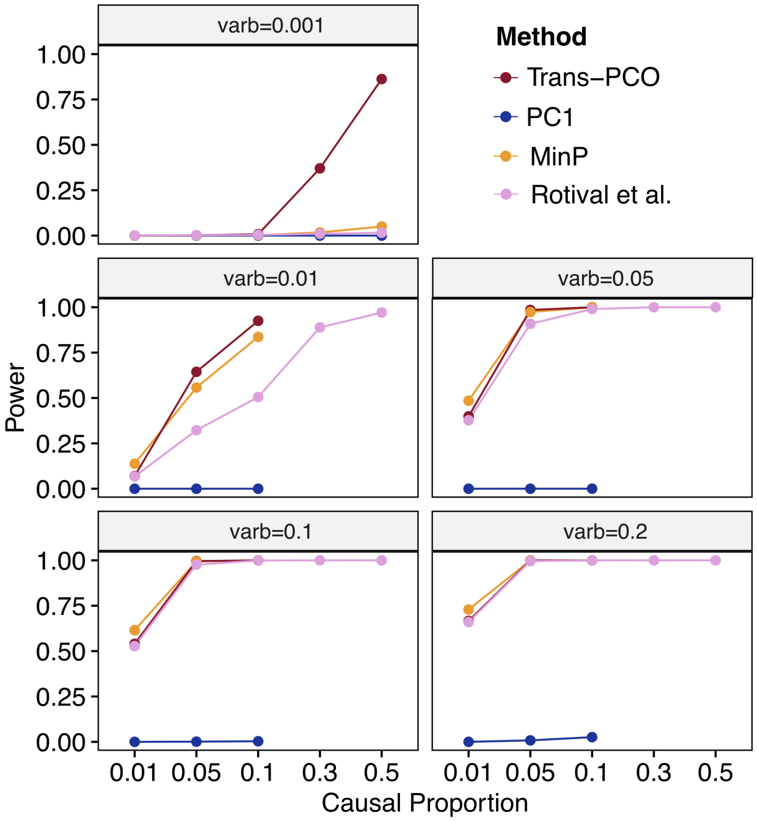


**Figure S20. Comparison of trans-PCO and Rotival et al.[10].** We compared the power of trans-PCO and the method proposed in Rotival et al. across various causal proportions under different genetic variances. Specifically, we simulated the proportion of target genes with non-zero effects to be (1) high levels of sparsity 1%, 5%, 10%, and (2) low levels of sparsity 30%, and 50%, and genetic variances to be (1) small effect 0.001, and (2) large effect 0.01, 0.05, 0.1, and 0.2. We used the same gene module as in Figure 2 and simulated the sample size to be 500. Rotival et al. method used a hypergeometric test to calculate enrichment p-values. P-values were corrected by ‘qvalue’ to control false positive rate at 10%. Power was computed from 10k SNPs across 1000 simulations.

### Supplementary Tables

Table S1. 166 co-expression gene modules in DGN

Table S2. Simulation results in Figure 2

Table S3. A summary of the number of reported signals

Table S4. 3899 significant *trans*-eSNP-modules pairs in DGN at 10% FDR

Table S5. Colocalization of *trans*-eQTLs with *cis*-eQTLs and *cis*-sQTLs in DGN

Table S6. Gene ontology enrichment of the nearest genes of *trans*-eQTL loci in DGN

Table S7. Functional annotations of 166 gene co-expression modules

Table S8. The list of complex traits and diseases used in colocalization analyses with *trans*-eQTLs

Table S9. The number and proportion of *trans*-eQTLs that colocalized with each complex trait and disease in Figure 5A

Table S10. The colocalization results of 179 *trans* loci with complex traits

Table S11. Heritability enrichment of blood traits in gene module 4 in Figure 5C

Table S12. 8116 (*trans*-eSNP, gene module) pairs detected by trans-PCO in eQTLGen

Table S13. 38 *trans*-eQTL signals in DGN that are replicated in eQTLGen

Table S14. Gene ontology enrichment of the nearest genes of *trans*-eQTLs loci in eQTLGen

Table S15. 50 MSigDB hallmark gene sets representing well-defined biological states and processes

Table S16. 965 significant *trans*-eSNP-modules pairs in DGN dataset with 50 MSigDB gene sets

Table S17. The colocalization results of *trans*-eQTL loci of 50 MSigDB hallmark gene sets with complex traits

Table S18. 2051 significant *trans*-eSNP-modules pairs for 11 MSigDB gene sets in eQTLGen

Table S19. Allergy drug target genes and their *trans* associated immune-related gene sets

Table S20. Enrichment of allergy drug targets in *trans* loci associated with immune-relevant gene sets
